# Exploring the Complex Dynamics of an Ion Channel Voltage Sensor Domain via Computation

**DOI:** 10.1101/108217

**Authors:** Lucie Delemotte, Marina A. Kasimova, Daniel Sigg, Michael L. Klein, Vincenzo Carnevale, Mounir Tarek

**Author notes:** Current address: Science for Life Laboratory, Department of Theoretical Physics, KTH, Box 1031, SE-171 21 Solna, Stockholm, Sweden.

## Abstract

Voltage-gated ion channels are ubiquitous proteins that orchestrate electrical signaling across excitable membranes. Key to their function is activation of the voltage sensor domain (VSD), a transmembrane four alpha-helix bundle that triggers channel opening. Modeling of currents from electrophysiology experiments yields a set of kinetic parameters for a given channel, but no direct molecular insight. Here we use molecular dynamics (MD) simulations to determine the free energy landscape of VSD activation and to, ultimately, predict the time evolution of the resulting gating currents. Our study provides the long-sought-for bridge between electrophysiology and microscopic molecular dynamics and confirms, as already suggested on the basis of experiments, that rate-limiting barriers play a critical role in activation kinetics.

## Introduction

Despite the complexity of the molecular processes underpinning electrical excitability of cells, the macroscopic currents measured in a typical electrophysiology experiment obey remarkably simple mathematical laws expressed in terms of few state variables. This observation, originating from the pioneering work of Hodgkin and Huxley, is replete with deep consequences: voltage gated ion channels (VGCs), the membrane proteins responsible for such currents, ought to cycle through a small number of distinct structural states. Accordingly, electrophysiology experiments are usually interpreted using the discrete state Markov (DSM) models approach, whereby a set of parameters characterizes each VGC under given conditions (*1*, *2*). This kinetic "discreteness" has been traditionally interpreted as the consequence of few well-defined free energy basins separated by rate-limiting free energy barriers. Identifying the structure associated to each of these minima is a long-standing scientific endeavor that has benefitted from the contribution of various approaches such as mutagenesis experiments (e.g. charge reversal and disulfide locking) (*3*–*6*), X-ray crystallography (*7*, *8*) and MD simulations (*9*–*12*). The significance of these studies that relate the electrical properties of VGCs to their physico-chemical determinants (*4*, *9*, *12*–*15*) is clear: a molecular level picture is key to understand the modulatory role of a myriad of physiological factors and to develop novel modulators.

Here, we focus on the mechanism controlling gating of VGCs, *i.e.* the transduction of a change in the transmembrane potential into a conformational transition involving the so-called voltage-sensor domain. The VSD is a structural domain of four transmembrane helices, from S1 to S4, that has been found thus far in VGCs (*16*), in voltage gated proton channels (Hvs) (*17*) and in a class of voltage-sensitive phosphatases (VSPs) (*18*).

The voltage sensing mechanism has been extensively studied and a consensus on the basics of the activation mechanism has been achieved: among the four helices, S4 plays a crucial role in voltage sensing. This helix carries several positively charged residues located inside the membrane and stabilized by the electrostatic interactions with the negative amino acids from the rest of the bundle. Upon application of a depolarizing electric potential, positively charged residues drag S4 across the membrane along the direction of the electric field, giving rise to a so-called gating current. The gating current reflects the total charge *Q* displaced and is a common reporter of the VSD conformational change that can be accessed experimentally. Finally, the motion of S4 opens the four-helices bundle allowing proton transport in Hvs, triggers catalytic activity in VSPs and opens the pore in VGCs.

Despite the fact that the sequence of events leading to activation of the VSD has become increasingly clear since the first experimental determination of a voltage-gated ion channel structure in 2005, many questions remain unanswered about the equilibrium populations of the different conformational states and the rate of exchange between them. In particular, an elusive question concerns the nature of the mechanical coupling between the VSD and the rest of the channel: to which extent can thermodynamics and kinetics of activation be described as intrinsic properties of the VSD? How do the coupled structural domains affect this process? A recent manuscript by Zhao and Blunck provided novel insight on this issue (*19*): in a breakthrough experimental investigation, the authors showed that the isolated VSD (iVSD) from Shaker is a voltage-sensitive cation channel with distinct thermodynamics and kinetics of activation (compared to the full-length channel, the voltage dependence of activation is less pronounced and the kinetics is slower). The iVSD construct is remarkable for a slow relaxation from a conducting to a non-conducting state. This mode shift into the “relaxed” state is associated with a hysteresis in voltage sensor activation curves when measured using activating versus deactivating voltage steps. Mode shifts of voltage sensor activation had been previously inferred from experiments using full-length channels (*20*–*22*) as well as a voltage-gated phosphatase (Ci-VSP) that is homologous to the VSD of Kv channels (*23*). The mechanism for relaxation remains poorly understood. In the case of iVSD, Zhao and Blunck provide evidence that the relaxed sensor is characterized by a single energetically favored state located at the end of the activation sequence (*19*). They argue that the relaxed state is kinetically distinct from earlier states through a relatively slow transition that may or may not be voltage-dependent. This differs from previous models in which mode shifting from the normal activation sequence involves a voltage-independent transition to a parallel sequence of relaxed states (*24*, *25*).

Here, we report the results of a computational investigation of the activation kinetics of an isolated VSD. We based our calculations on a model of the sensor of Kv1.2, a close relative of Shaker for which a crystal structure of the activated state is available. Specifically, we have calculated potentials of mean force (PMFs, or free energy landscapes) from several microsecond-long all-atom atomistic molecular dynamics simulations, thereby broadening the scope of a previous investigation by Rudy *et al.* (*14*). We have then calculated the associated gating currents and compared them to electrophysiology results. The good agreement between MD-derived gating currents and experiment lends confidence in our approach and thus, ultimately, in the molecular picture of the mode shift effect that we obtain from our calculations. Significantly, our strategy to bridge the gap between microscopic and macroscopic dynamics of ion channels is completely general and transferable. This study paves the way for a wealth of future investigations aimed at understanding the kinetic consequences of modulatory factors like drugs or lipids.

## Results and discussion

### Modeling the structural states of Kv1.2 VSD

Experimental and simulation studies have revealed the salient features of the VSD motions associated to Kv1.2 gating: overall, four sequential transitions bridge five separate “conformational states”, called ε, δ, γ, β and α. The latter are defined in terms of specific patterns of salt bridges between the mobile positive (gating) charges and the quasi-static negative charges within the VSD and at the water/membrane interface (*4*, *9*, *12*) (Figure 1A). In particular, there are six positive charges on S4 (R294 (R1), R297 (R2), R300 (R3), R303 (R4), K306 (K5) and R309 (R6)), two negative charges on S2 (E226 and E236), one on S1 (E183), and one on S3 (D259). In the fully activated α state, the S4 helix is in the most “upward” position: R1 and R2 interact with the negatively charged moieties of the lipid headgroups of the upper leaflet, R3 and R4 are in contact with the two outermost negative charges, E183 and E226, and K5 and R6 form salt bridges with the two innermost negative charges, D259 and E236. In the intermediate states, from β to δ, the positively charged residues are sequentially shifted by a one-turn-of-a-helix step downward across the membrane, switching their countercharges to the closest negatively charged residues or lipid headgroups (Figure 1A, B). Finally, in the fully resting ε state, the S4 helix is in its most “downward” position: R1 and R2 interact with D259 and E236, and R3, R4, K5 and R6 all form contacts with the lipid headgroups of the lower leaflet (Figure 1A).

**Figure 1.**
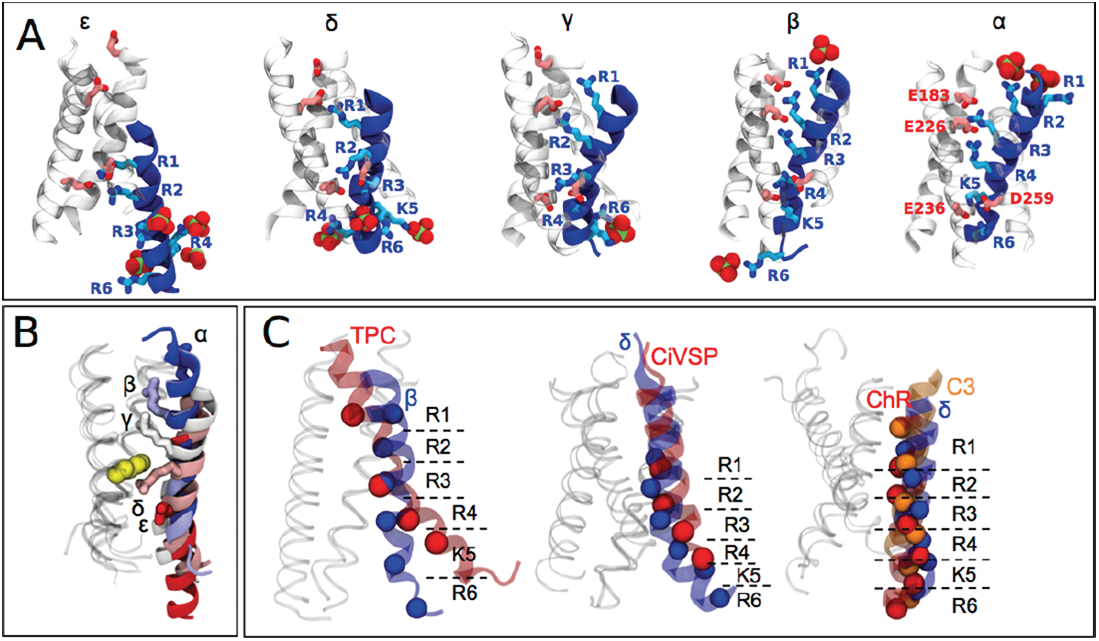
A) Starting structures representing the five activation states of the Kv1.2 VSD. From the resting (ε) to activated (α) conformations, the S4 helix (blue) moves upwards. The shift between the activation states involves a concerted rearrangement of the salt bridges between the S4 positive charges, from R1 to R6 (cyan sticks), and the negative charges of S1-S3 (pink sticks) and the lipid headgroups (red spheres). B) Superimposition of the five starting structures of the activation states; each structure is roto-translated to optimally superimpose the S1-S3 segments. S1-S3 and S4 are respectively shown as transparent ribbons or as a solid ribbon of colors ranging from red (ε) to blue (α). R2 is represented as sticks (the color, from red (ε) to blue (α), depends on the activation state), and the gating charge transfer center F233 is shown as yellow spheres. Note how the ratchet-like movement of S4 brings R2 from an inner (ε) position to an outer (α) one. C) Comparison of the starting structures with the available resting-state structures and the resting-state models. The position of S4 in the β state resembles closely that of TPC1 [PDB ID 5E1J (*26*)], while this position in the δ state is similar to Ci-VSP [PDB ID 4G80 (*27*)] and the structural models by Jensen et al. (Chr) (*12*) and Henrion et al. (*4*) (C3), respectively. The spheres represent the Cα atoms of the R1, R2, R3, R4, K5 and R6 residues.

Among these five states of Kv1.2, the fully activated one (α) has been resolved by X-ray crystallography (*28*) while the structures of ε, δ, γ and β are not available yet. To construct the molecular models of these states, we applied the following strategy: first, we aligned the sequence of the α Kv1.2 VSD with the ones of the other states, shifting the S4 segment of the target sequence toward the N-terminus (by one, two, three or four helical turns for β, γ, δ and ε respectively); then, we generated homology models of β, γ, δ and ε using the structure of the α state as a template (Figure 1A, B); finally, to verify that these molecular models indeed reflect voltage-driven conformational rearrangements in the VSD, we compared them with the available structures of resting states of VSD-containing proteins (TPC1 (*26*) and CiVSP (*27*)) and with molecular models of Shaker-like channels published previously (*4*, *12*) (Figure 1C). In particular, we found that the β and δ states resemble closely the structure of the resting state VSDs in TPC1 and Ci-VSP, respectively (Figure 1C). These similarities and the notion of a conserved voltage-sensing mechanism across different voltage-sensing protein families (*16*) gives us confidence in the models of the intermediate structural states of Kv1.2 we generated. Also, the δ state aligns with other molecular models of the resting states that were either built using experimental constraints (*4*) or obtained through multi-microsecond MD simulations under hyperpolarization conditions (*12*) (Figure 1C). The fully resting ε state does not have a homolog in the available structural data, and indeed its existence is debated (*10*). Our calculations serve also the purpose of assessing its stability and accessibility under physiological conditions.

### Free energy landscape of VSD activation

To monitor the progression along the activation pathway, we have characterized the free energy landscapes for each of the four transitions. To do so, we have designed a set of geometric descriptors (or collective variables - CVs) (*29*), that take into account the two major molecular determinants of VSD activation, as determined in Delemotte et al. (*9*): (a) the unbinding [binding] of the gating charge (Ri) from [to] the initial [final] binding site and (b) the spatial translation of the gating charges along the distance vector between two binding sites. Indeed, taking advantage of the spatial proximity of several negatively charged groups of the VSD, the number of binding sites can be reduced to four: (*Bi*) phosphate groups of the top lipid layer; (*Bii*) top protein binding site comprising E183 and E226; (*Biii*) bottom protein binding site comprising E236 and D259; and (*Biv*) phosphate groups of the bottom lipid bilayer (Figure 1 – Figure supplement 1). The two bottommost transitions (ε/δ and δ/γ) are effectively described by two collective variables. In particular, the ε/δ transition is described by *CV*_*R1*_ (transfer of R1 from binding sites *Biii* to *Bii*) and *CV*_*R3*_ (transfer of R3 from *Biv* to *Biii*). The δ/γ transition, on the other hand, is described by *CV*_*R2*_ (transfer of R2 from *Biii* to *Bii*) and *CV*_*R4*_ (transfer of R4 from *Biv* to *Biii*). Because of the involvement of the binding site *Bi* in the activation mechanism, the two uppermost transitions (γ*/*β and β*/*α) require three CVs. The γ/β transition is described by *CV*_*R1*_ (transfer of R1 from *Bii* to *Bi*), *CV*_*R3*_ (transfer of R3 from *Biii* to *Bii*) and *CV*_*K5*_ (transfer of K5 from *Biv* to *Biii*). The β*/*α transition is described by *CV*_*R2*_ (transfer of R2 from *Bii* to *Bi*), *CV*_*R4*_ (transfer of R4 from *Biii* to *Bii*) and *CV*_*R6*_ (transfer of R6 from *Biv* to *Biii*).

We have thus used multiple-walkers well-tempered metadynamics to build multidimensional PMFs (free energy landscapes) for each of the four transitions (Figure 2A). Adaptive enhanced sampling methods like metadynamics have the advantage of allowing repeated exploration of the same regions of the configuration space; consistent estimation of the free energy can be thus used as criterion for convergence and to calculate statistical errors (Figure 2 – Figure supplement 1). In particular, this approach allows one to ascertain that the CVs used to characterize the PMF (and along which sampling is enhanced) indeed encompass the slowest degrees of freedom of the process (*30*). A total of 73 µs aggregated simulation data allowed us to explore the conformational landscape well beyond the initial conformations (labeled as stars in Figure 2A). Of particular note is that the starting conformations did not always correspond to free energy minima. However, we could sample several free energy minima located in their neighborhoods thanks to the use of this adaptive enhanced sampling method.

**Figure 2.**
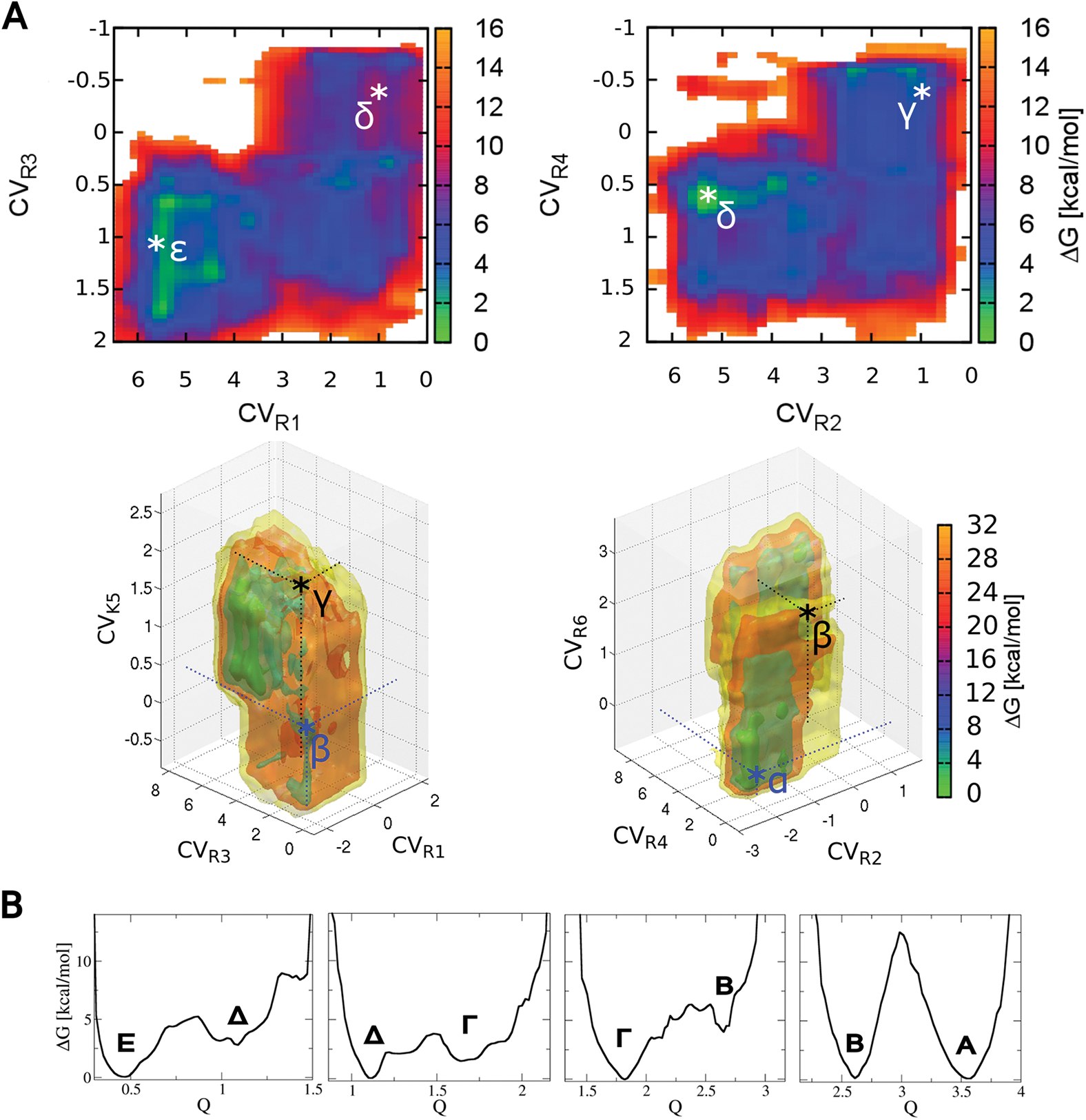
A) PMFs (Free energy landscapes) of the 4 corresponding transitions in two- (ε/δ and δ/γ transitions) and three- (γ/β and β/α transitions) dimensional CV space estimated from the metadynamics runs. Regions of low and high free energy are depicted in cold (green to blue) and in hot colors (red to orange), respectively. The CVs of the starting configurations are highlighted with the * symbol. B) Reweighted PMFs along the gating charge Q. Starting with the geometrical definition of the putative activation states ε to α, we have obtained larger ensembles of structures corresponding to stable thermodynamic states E to A. All free energies are reported in kcal/mol, and Q in electronic units (eu).

We then used the sampled VSD configurations to calculate the PMF as a function of the experimental observable gating charge *Q*, i.e. the time integral of the gating current (Figure 2B). We find that the free energy landscape along *Q* is described by a series of metastable states and that one barrier dominates the landscape (Figure 2B). The ensembles of configurations relative to each free energy well correspond, albeit loosely, to the states ε to α. This is strong evidence that the intermediate configurations uncovered by previous experimental and simulations studies are in fact thermodynamically metastable. Hereafter, we refer to these states as Ε, Δ, Γ, Β and Α. The expected values of Δ*Q* (transition gating charges) for the four transitions in the activation pathway are 0.62, 0.69, 0.84, and 0.93 electronic units (eu), respectively. The barriers between the Ε, Δ*, Γ* and Β states are relatively small (<5.5 kcal/mol) and larger in the forward direction (from Ε to Β) (activation) than in the reverse one (deactivation). Thus according to our results, the E state is accessible under physiological conditions and is a part of the activation sequence. The final barrier between Β and Α stands out as being more than twice as high as the next largest barrier that precedes it, *i.e.* ~ 12.5 kcal/mol, the two states being of similar stability. The analysis of the number of crossing events for each transition (Table 1) reveals that the transitions were not sampled enough to have confidence in the quantitative estimation of the free energy barriers; however, given that the number of barrier crossings is similar in all of the transitions, we are confident that the Β/Α transition is significantly higher compared to the others.

**Table 1:** Statistics of barrier crossing events during the metadynamics trajectories.^c^.

|  |  | # initiated | # entering the | # reaching | # full |
| --- | --- | --- | --- | --- | --- |
| Δ-E | $\Delta$ | 21 | N/A<sup>a</sup> | 3 | 1 |
|  | E | 21 | N/A<sup>a</sup> | 7 | 1 |
| Γ-Δ | $\Gamma$ | 21 | N/A<sup>a</sup> | 6 | 1 |
| | $\Delta$ | 21 | N/A<sup>a</sup> | 5 | 0 |
| B-Γ | B | 21 | 4 | N/A<sup>b</sup> | 1 |
| | $\Gamma$ | 21 | 21 | N/A<sup>b</sup> | 5 |
| A-B | A | 21 | 9 | 5 | 0 |
|  | B | 21 | 9 | 3 | 1 |

<sup>a</sup>N/A stands for not available. In the $\Gamma$ - $\Delta$ and the $\Delta$ -E transitions, the barrier region is not well defined, whereas the barrier itself is defined ( $Q=1.50$ e and $Q=0.75$ e, respectively).
<sup>b</sup>In the B- $\Gamma$ transition, the precise placement of the barrier is not well defined, whereas the barrier region is defined ( $2.00e < Q < 2.45e$ ).
<sup>c</sup>For each of the transitions, 21 walkers were initiated in each basin. We report the number of walkers initiated in each basin that enter the barrier region, the number of walkers that reached the top of the free energy barrier and the number of walkers that reach the other basin of the transition (labeled as full crossing).
Note also that our previous estimation of the $\Delta$ /E free energy landscape led us to a qualitatively similar picture, with an activation barrier of similar height (29). However, the E and $\Delta$ states had previously been found to be of similar free energy. In the present paper, we have more than doubled the sampling (with a total of more than 10 μs simulation time for the sole $\Delta$ /E transition) and applied a more robust reweighing procedure (see Methods), which makes the present estimation more reliable.

Note also that our previous estimation of the Δ/E free energy landscape led us to a qualitatively similar picture, with an activation barrier of similar height (*29*). However, the E and Δ states had previously been found to be of similar free energy. In the present paper, we have more than doubled the sampling (with a total of more than 10 µs simulation time for the sole Δ/E transition) and applied a more robust reweighing procedure (see Methods), which makes the present estimation more reliable.

### Mechanism of VSD activation

The metadynamics calculations allowed us to identify the conformational ensembles corresponding to the different activation (E to A) and transition states. The representatives of these conformational ensembles (nine in total, Figure 3) were selected as the structure with the lowest RMSD with respect to all the other configurations in the ensemble. The activation states resemble closely the starting configurations; in particular, the pattern of the salt bridges between S4 and the counterpart negative charges is identical. The main feature of the transition states is the cation-π interaction of the positive charge moving between *Biii* to *Bii* with F233, a residue that is conserved in all voltage sensors characterized so far. The crucial role of this residue in the translocation of positive charges along the direction of the electric field was previously highlighted using mutagenesis experiments (*3*). Due to its specific position in the voltage sensor, F233 separates the upper and lower water crevices of the four-helix bundle, constituting the so-called hydrophobic plug. This unique configuration focuses the electric field in the center of the voltage sensor and, in particular, on the positive charge moving between *Biii* to *Bii* (*9*). Based on its role in the voltage sensing mechanism, F233 is accordingly called the gating charge transfer center.

A novel and so far unreported feature that our calculations reveal is a change in the pattern of interactions between the extracellular linkers, S1-S2 and S3-S4, along the activation process (Figure 3 – supplement 1, highlighted with white circles). In particular, while in the most activated states (B and A), these linkers establish a network of interactions, in the others (E, Δ and Γ), S1-S2 and S3-S4 are not in contact with each other. Moreover, during the transition between the B and A states, the set of interacting residues of the S1-S2 linker changes from 206-210 to 211-216. Interestingly, in the molecular dynamics trajectories previously obtained for the Kv1.2/2.1 paddle chimera subject to a hyperpolarized potential (*12*), a significant conformational rearrangement of the S1-S2 and S3-S4 linkers also takes place before an abrupt switch from the most activated state to the next one in the sequence of deactivation. The conformational change of the voltage sensor extracellular linkers is therefore seemingly an integral part of the B/A transition and might be the origin of the large free energy barrier.

**Figure 3.**
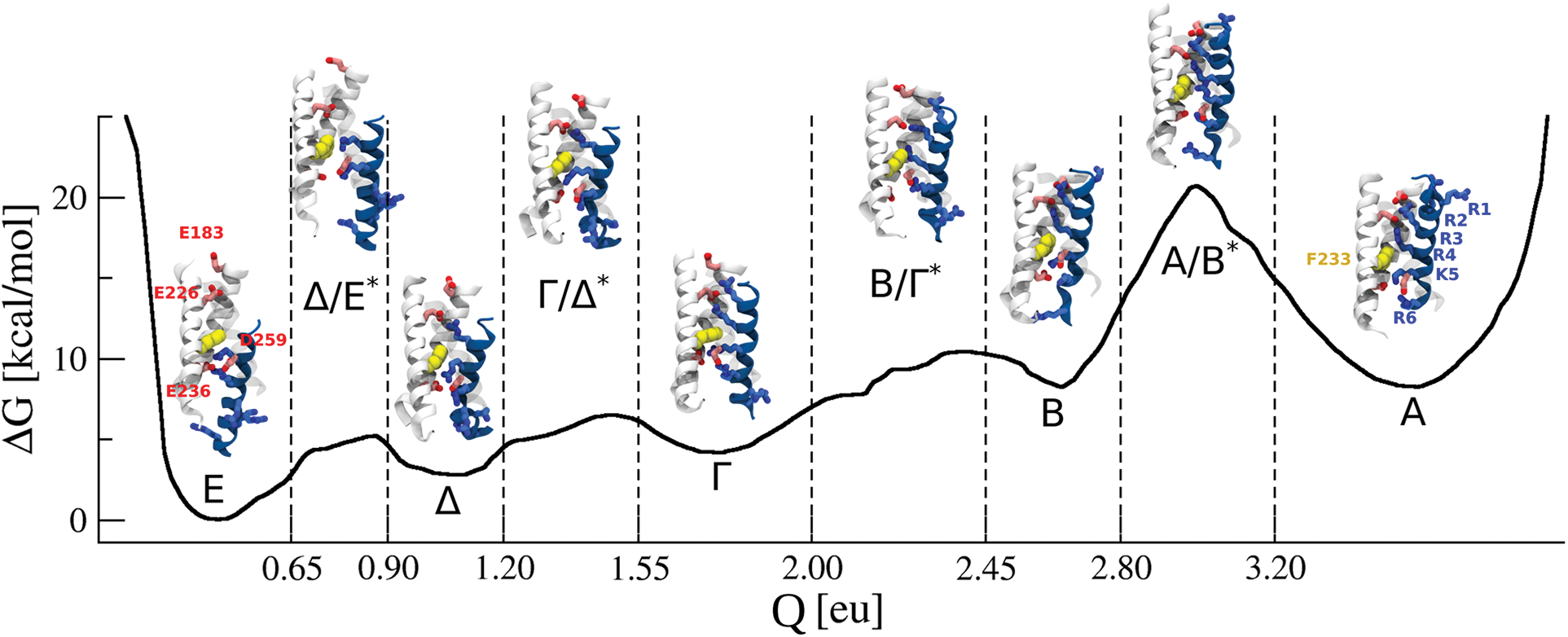
Representative configurations of the five free energy minima and four transition states, defined as the structures with the lowest RMSD with respect to all the other configurations in the ensemble. The minima and barrier ensembles are defined based on the values of the gating charge (horizontal axis). The colors and representations of the molecular models are the same as in Figure 1.

A prominent secondary structure changes can be seen in the S4 helix during deactivation. In the A state, the bottom portion of S4 fluctuates between α- and 3_10-_ helices. Transitioning to the B state, however, this segment adapts mostly a 3_10_ helix and rarely an unfolded conformation. Finally, in the bottommost activation states, this portion of S4 corresponds to an α-helix. Interestingly, a 3_10_ conformation appears in some of the configurations in the middle and upper portion of S4 during the transitions from Γ to Ε. This propensity of the S4 section traversing the gating charge transfer center for the 3_10_ conformation, although only present in a fraction of instantaneous configurations, is in qualitative agreement with the results obtained previously (*4*, *31*). We also note that the S3-S4 linker, which is mostly in an α-helical conformation in the bottommost states of the activation sequence (from Δ to E) and in the fully activated state (A), has a tendency to get unstructured in the B state. Strikingly, in the Γ state the S3-S4 linker and S4 form a continuous α-helix.

### In silico electrophysiology

In order to compute the gating currents of the Kv1.2 voltage sensor and to compare them to the available experimental data, we reconstructed the free energy landscape *G(q)* underlying the activation process. To do so, we combined the four PMFs obtained separately for each transition into a continuous profile; the result of this is presented in Figure 4. Contrary to expectations from electrophysiology recordings on Shaker-like channels, the resting ε state has a lower free energy than α. The origin of this inconsistency remains to be explained.

**Figure 4.**
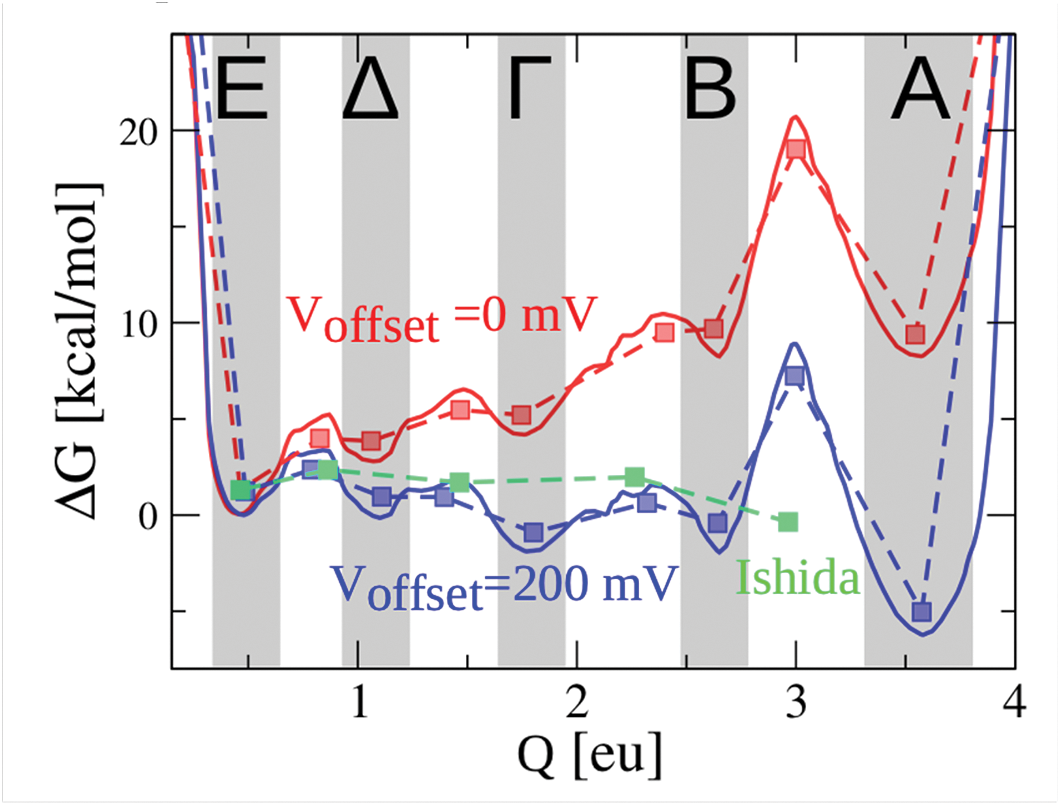
Ab initio PMF of the 5-state activation sequence obtained by combining the four PMFs shown in Figure 2B (solid red line). The landscape depicted in blue is the same free energy profile after applying a +200 mV offset. The coarse-grained discrete state Markov (DSM) positions of wells and barriers are indicated by squares of the same color. For comparison, the DSM landscape of an individual subunit in the Ishida et al. model (*33*) is shown in green.

In order to better compare the voltage-dependence of simulated gating currents with experimental data, we removed the upward tilt of the landscape through a simple voltage offset. As a basis for comparison, we considered the Q-V curve derived from the 3-state activation sequence in an individual voltage-sensing subunit in the full-channel gating model of Kv1.2 suggested by Ishida et al. (*33*). A +200 mV voltage offset was required to bring about a rough alignment between the partial Ishida model and the 5-state discrete state Markov (DSM) model obtained by coarse-graining the continuous PMF (*1*) (Figure 4). Note that in the 3-state scheme by Ishida et al., the gating charge transfer (2.5 eu) is about 16% greater than the electrical distance from E to B states in the PMF profile (2.1 eu); however, it is less than the total distance from E to A (3.1 eu) (Figure 4).

An important experimental quantity is the equilibrium position of charge as a function of voltage (Q-V curve). The Q-V may be derived from the voltage dependent PMF *W*(*Q*,*V*) = *G*(*Q*) – *QV* through ∫ *p*_*eq*_ *QdQ*, where 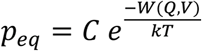 is the Boltzmann equilibrium probability with *C* as the normalization constant. Experimentally, Q-V curves are typically obtained by integrating a series of gating currents to different voltages starting from the same holding potential *V*_*h*_. If there are long or short time scales of gating charge transfer that cannot be captured by experiment, the measured Q-V curves may show a dependency on *V*_*h*_. This mode shift effect seen as a hysteresis in the Q-V is a common feature among voltage sensors, including the recently characterized iVSD (*19*). Figure 5 features computed Q-V curves predicted by the PMF. The profile of the PMF derived in this study suggests there may be a substantial time scale difference between transitions across the dominant free energy barrier, Β/Α, and those traversing the smaller barriers between the Ε and Β states. The experimental quantity describing the rate of *Q* transfer is the gating current. We should expect short time-scale gating currents from a negative *V*_*h*_ to be reflected in the Q-V curve obtained from the abbreviated landscape E-B. On the other hand, long time-scale gating currents obtained from an arbitrary choice of *V*_*h*_ will include transitions across the Β/Α barrier, whose inclusion in the PMF landscape generates a steeper and more left-shifted Q-V (Fig. 5). The question of which Q-V is the operational one for the time scale of channel activation depends on the value of the effective diffusion coefficient *D* for gating charge movement.

**Figure 5.**
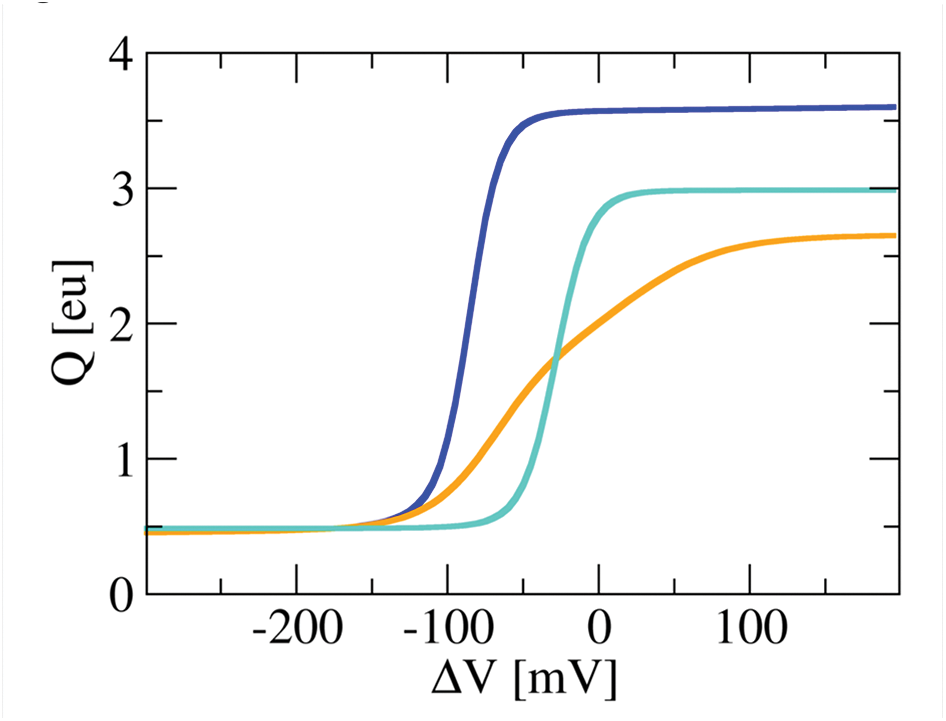
Gating charge as a function of voltage (Q-V). The Q-V relationships are calculated from the 4-state E to B (orange) and 5-state E to A (blue) PMFs. For comparison, the 3-state DSM Ishida et al. Q/V curve is shown in cyan. Note that the 4-state Q-V is more shallow and right-shifted compared to the 5-state version.

Computing MD-derived gating currents requires solving the time evolution of the probability distribution *p*(*Q*,*V*,*t*) of *Q* as it undergoes Brownian motion across the calculated PMF. We assume the dynamics is governed by the Smoluchowski equation:

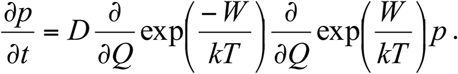

Initial conditions corresponding to the resting potential *V*_*r*_ were determined from the Boltzmann equilibrium distribution as described above. After a step change in voltage, the system relaxes to a new equilibrium as determined by the Smoluchowski equation. The mean gating current for this process was computed as 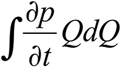.

To follow a truly *ab initio* approach, we estimated *D* using molecular dynamics simulations. To do so, we split the entire *Q* space into 0.2 e bins and initiated molecular dynamics simulations in each of them (Figure 6). Note that we modified the interatomic potential by adding a bias obtained previously from the metadynamics simulations; by doing this we were able to flatten the free energy landscape and therefore deal with a purely diffusive system. We then calculated *D* by estimating mean square displacement (MSD) of the collective variable *Q* as a function of time. We found that *D* is very similar for all the transitions and corresponds to 180±110 eu^2^m^−1^ for E/Δ, 290±130 eu^2^m^−1^ for Δ/Γ, 150±50 eu^2^m^−1^ for Γ/B, and 330±230 eu^2^m^−1^ for B/A (Figure 6). These values are, however, too large to compute gating currents compatible with the experimental data: for comparison, the diffusion coefficient estimated from measurements of fast gating current transients in the Shaker channel is 41 eu^2^/ms, i.e. about one order of magnitude smaller (*34*). Therefore, we adopted an empirical approach and chose a value of *D* that was roughly consistent with the time course of published gating currents.

**Figure 6.**
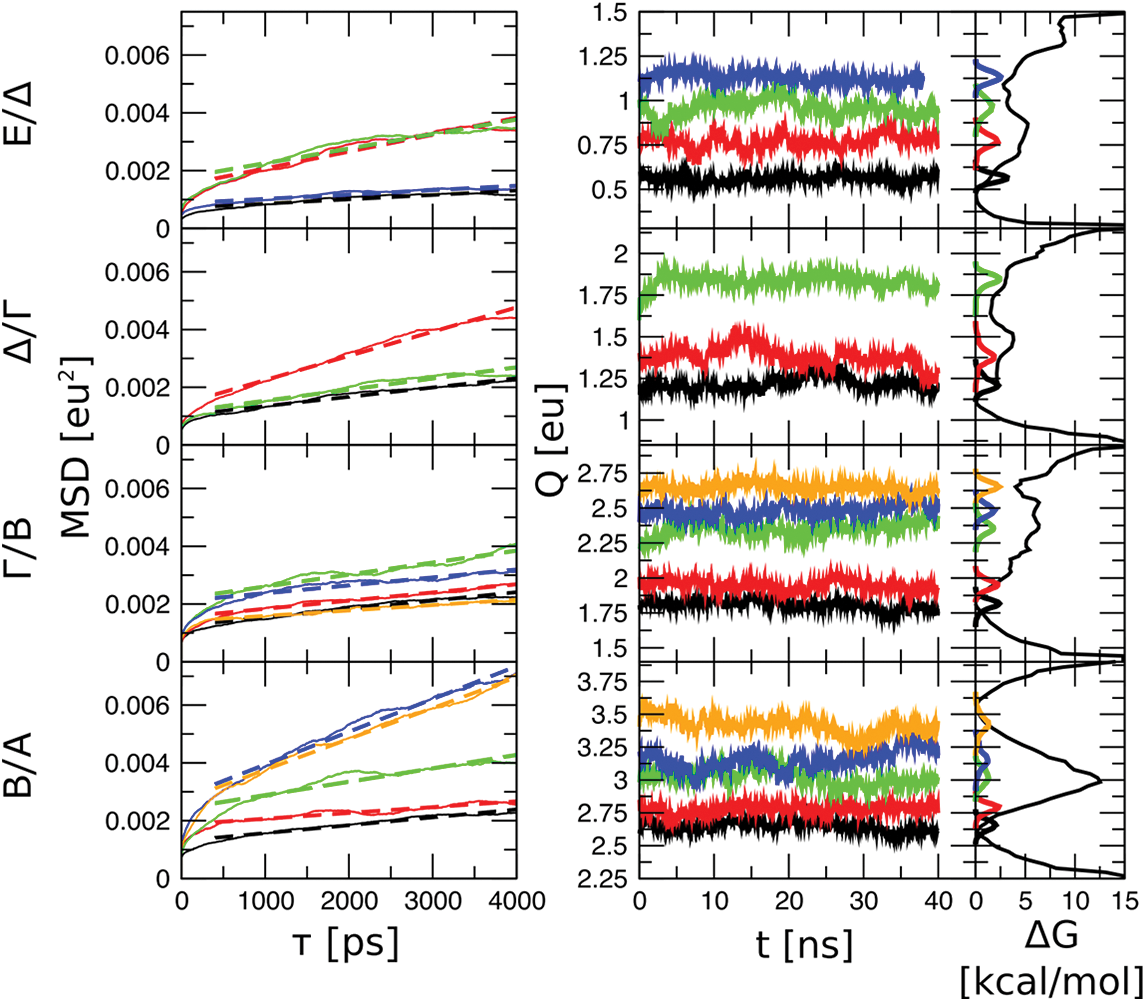
ab initio calculation of the diffusion coefficient in the four voltage-sensor domain transitions (E/Δ, Δ/Γ, Γ/B and B/A) obtained via biased MD simulations. Left column: mean square displacement (MSD) of the gating charge Q as a function of time (solid lines). Black, red, green, blue and orange lines correspond to the different replicas with distinct initial values of Q (see the middle column). The diffusion coefficient was estimated as half of the slope of the best linear fit between 400 and 4000 ps (dashed lines): for E/Δ, 180±110 eu^2^m^−1^; for Δ/Γ 290±130 eu^2^m^−1^; for Γ/B, 150±50 eu^2^m^−1^; for B/A, 330±230 eu^2^m^−1^. Middle column: time series of Q for all the replicas considered in the diffusion estimation. Right column: the distributions of Q are overlaid to the PMF showing that the major basins and barriers have been sampled during the runs.

From the +200 mV offset PMF and a constant value of *D* (to be determined), we computed gating currents by solving the Smoluchowski equation using two methods (*35*). In the first method, the Smoluchowski equation was cast as a stochastic differential (Langevin) equation and *Q* trajectories were simulated (Figure 7A). Gating current trajectories (Figure 7B) were obtained by applying a low-pass filter to the *Q* trajectories, resampling to the time scale of interest, and then computing the time derivative of the filtered traces. The result is a series of spikes resembling filtered shot noise (*35*). An ensemble of individual traces can be averaged to obtain the mean gating current, though this is inefficient from a computational standpoint. The preferred method involved discretizing the Smoluchowski equation into a master equation in the form of a 400 x 400 tridiagonal matrix. This matrix was symmetrized and eigenvector analysis yielded the relaxation time constants and amplitudes for each exponential component of the mean gating current (Figure 7C). The sum of exponents were then subject to low-pass filtering. A third method involves the systematic coarse-graining of the energy landscape into discrete state and barrier points (Figure 4), yielding Arrhenius-like rate constants with *D* as the prefactor (*1*). This last method is easily implemented and recapitulates the practical approach to data analysis using a master equation with a small number of states (DSM approach) but lacks information about fast rearrangements within states. These fast transient events (see early spike in the black trace of Figure 7C) are not typically of interest unless there is a merging of fast and slow time scales brought about by energy barriers of insufficient height, in which case coarse-graining fails and the transitions become diffusion limited and Markovian kinetics no longer apply.

**Figure 7.**
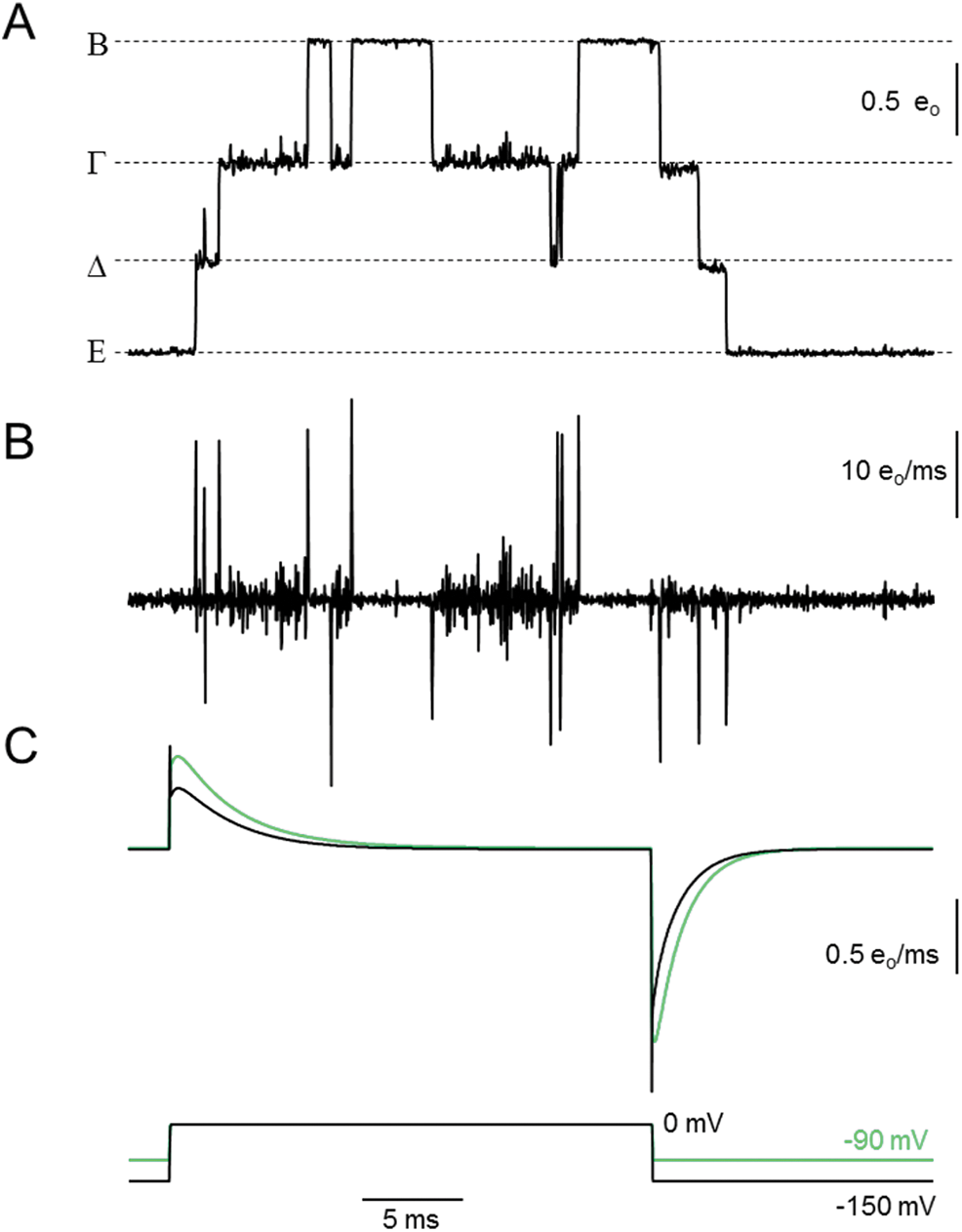
Gating charge kinetics calculated from the PMF with +200 mV voltage offset and diffusion coefficient D = 3 e^2^/ms. (A) ON and OFF Brownian motion trajectory of gating charge Q in response to a test pulse to 0 mV from a holding potential V_h_ = −150 mV. The voltage protocol is shown at the bottom of the figure. Simulation parameters were as follows: temperature = 297 K; time step for Langevin dynamics = 20 ns; Gaussian filter cut-off frequency = 10 kHz; digital sampling rate = 50 kHz; (B) Gating current trajectory corresponding to (A). (C) Eigenvector calculations of mean gating currents from the PMF landscape (black line, V_h_ −150 mV) and from the 3-state scheme by Ishida et al. (green line, V_h_ −90 mV).

The eigenvalue method was used to generate mean gating currents and this was compared to the predicted gating current of the Ishida et al. 3-state model (*33*) (Figure 7C). A value of *D* = 3 eu^2^/ms achieved the best overall match to the kinetics of the reference model. Note that the obtained mean gating current features a prominent rising phase, which is also seen experimentally and represents a common feature of voltage-gated ion channels. Using the same value of *D*, gating charge trajectories at 0 mV fluctuate among states E-B (Figure 7B) with long dwell times (compared to the actual time to cross barriers), consistent with a Markov process. The last state in the sequence, the Α state, is rarely visited despite being lower in free energy (Figure 7 – figure supplement 1). Thus, short-time experiments should be governed by the abbreviated 4-state scheme, leading to the shallow Q-V curve in Figure 5.

The relaxation time constants for the complete PMF are shown in Figure 8. We found that the slowest time constant pertaining to the Β/Α transition is clearly separated from the others, introducing two radically different time scales. In order that the Β/Α transition be included in the normal activation pathway, either the value of *D* would need to be seven orders of magnitude greater, or the B/A free energy barrier would need to be smaller than we calculated, perhaps as a result of interactions with the pore. The VSD-pore interaction occurs in late stages of the gating process according to current gating models (*37*). Time constants computed from the coarse-grained DSM representation in Figure 4 are shown as dashed curves. These are derived from rate constants and gating charge displacements evaluated at 0 mV, and extrapolated to a wider range of voltages using the standard Arrhenius form of the rate constant. It is apparent that beyond the voltage range of approximately −100 mV to +100 mV, the PMF-derived time constants deviate from the asymptotically linear DSM time constants as a result of diffusion-limited transitions caused by flattening of free energy barriers. This has implications for the traditional form of modeling ion channel kinetics based on the DSM paradigm; in particular, it challenges the usual assumption that the voltage dependence of decay constants is asymptotically exponential.

**Figure 8.**
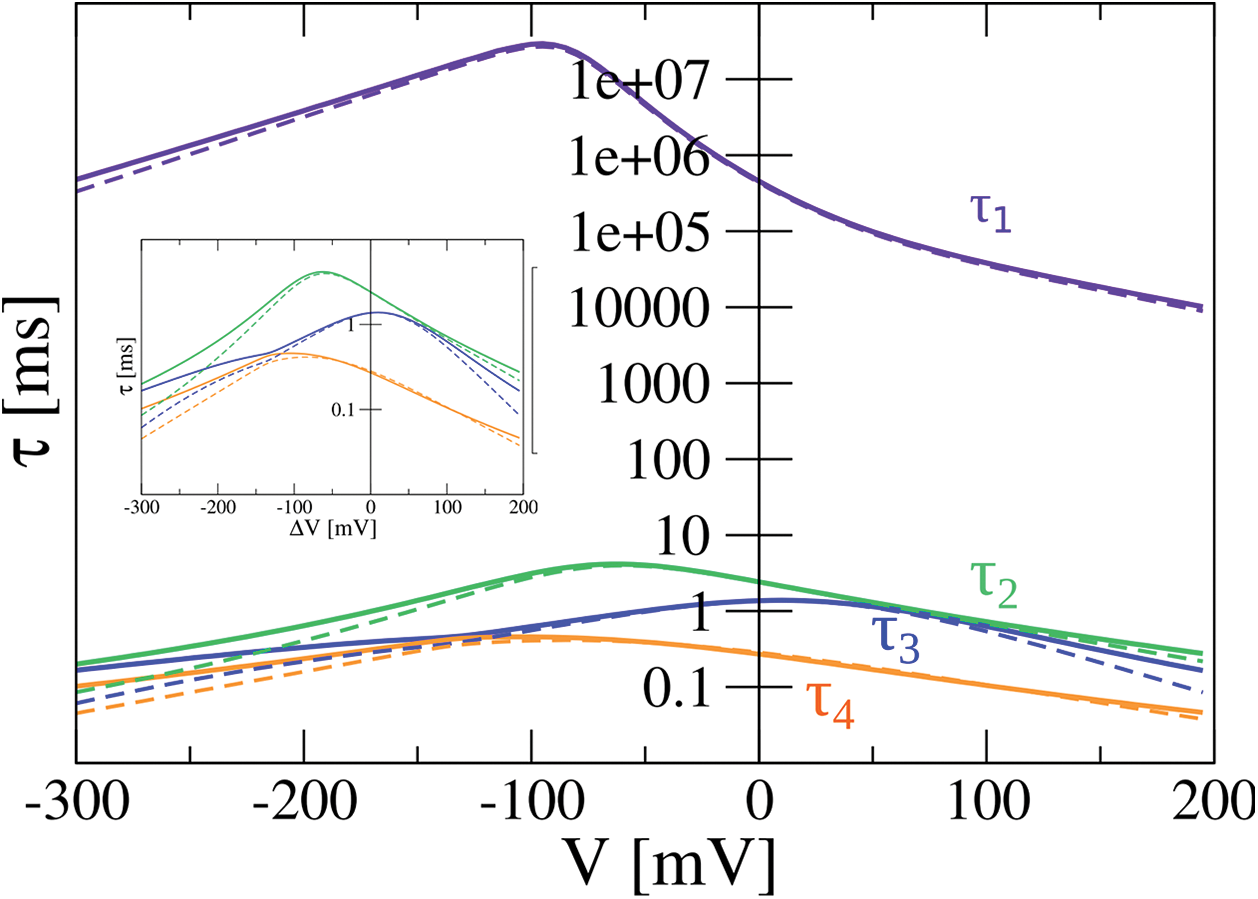
Slow decay constants (τ) of the four transitions versus voltage computed from eigenvalue analysis of the 5-state PMF profile (solid lines) and the coarse-grained DSM scheme (dashed lines). The inset displays a blown-up version of the three faster components.

## Conclusion

Our calculation of the voltage sensor activation free energy landscape represents an important milestone toward a quantitative, molecular-level, picture of an ion channel gating. The landscape along the gating charge reaction coordinate reveals several important features. First, five distinct metastable states are present along the activation sequence. Second, the four earlier states (E-B) are separated by relatively low (5 kcal/mol) free energy barriers whereas a large barrier has to be crossed to reach the last state (A). The free energy barriers (Δ*G*^*^) in the Ε-Β PMF transitions are substantially smaller than barrier enthalpies (Δ*H*^*^) in early gating transitions measured from temperature studies in the related Shaker channel (~5 kcal/mol versus ~15 kcal/mol) (*38*); the inference of a large entropy of activation (*T*Δ*S*^*^ = Δ*H*^*^ − Δ*G*^*^ ~ 10 kcal/mol) facilitating these early events warrants further study. The free energy landscape predicts two radically different timescales for the gating currents: states E-B equilibrate on the ms timescale while the A state is accessible only on the multisecond time scale. It remains to be seen whether VSD-pore interactions lower the B/A transition barrier.

The MD simulations presented here yield mechanistic insight into voltage sensor activation. First, our analysis confirms the presence of a previously described transient cation-π interaction between the positive charge traversing the electric field and the gating charge transfer center residue F233 in each of the transition states. Second, we have related the high free energy barrier to a restructuring of the S1-S2/S3-S4 linker interaction during the transition between states B and A. Support for this comes from extended MD simulations by Jensen et al. showing a sudden movement of S4 occurring after a long dwell time in the A state, but shortly after S1-S2/S3-S4 linker restructuring, followed by a fast deactivation process. Finally, as reported earlier, the S4 helix section that is traversing the gating charge transfer center has a tendency to assume a 3_10_ conformation.

Mode shifting has recently been shown to be an intrinsic feature of the voltage sensor domain activation, even in isolation from the pore domain. This implies the existence of one or more additional conformational states, sometimes referred to as being “relaxed”, that are visited only under prolonged depolarization conditions. We propose several lines of evidence that this relaxed state corresponds to the A state, which is separated from the regular activation states by a high free energy barrier. First, the predicted slowing of deactivation and the leftward shift of the Q-V curve achieved experimentally by starting from a *V*_*h*_ of 0 mV (corresponding at a molecular level to transitions from the A state) is also seen experimentally in the isolated iVSD construct (*19*). Second, gating currents predicted from the abbreviated activation sequence E-B demonstrate a characteristic rising phase that would be missing if the value of the diffusion coefficient were adjusted upwards to bring the B/A transition into the experimental time scale. Finally, Priest et al. reported that shortening the S3-S4 linker hinders deactivation from the relaxed state (*39*). This supports our observations that S1-S2 and S3-S4 linker interactions help shape the B/A transition barrier.

Future work will need to take into account the interactions between the VSD and the pore domain in order to fully evaluate the influence of the pore on the PMF and in particular on the Β/Α free energy barrier.

## Materials and methods

### VSD models generation

To produce a structural model for each activation state of the Kv1.2 VSD (besides the one solved in the crystal structure (*40*)) we threaded the sequence through the S4 structure by shifting the register (compared to the activated state) by three, six, nine or twelve residues (corresponding to one, two, three or four helical turns, respectively). The resulting structures were used as initial models for the conformational states along the deactivation pathway: β, γ, δ and ε, respectively, using the nomenclature proposed in previous work(*9*). We used MODELLER to generate these structures (*41*).

### System preparation

The ε, δ, γ, β and α state models of the Kv1.2 VSD (residues 163 (S1) to 324 (S4-S5 linker)) were then inserted in a fully hydrated 110x110 Å^2^ 1-palmitoyl-2-oleoyl-sn-glycero-3-phosphocholine (POPC) bilayer using the VMD membrane builder (*32*). Overlapping lipids and water molecules were then deleted. The system was then solvated with 150 mM NaCl. The total number of atoms in the five systems is ~ 100,000.

### Molecular dynamics simulations

The systems were equilibrated under normal constant temperature and pressure conditions (300 K, 1 atm.). The lipid tails were melted during the first nanosecond, restraining the position of the protein and lipid head groups to their initial position with a strong harmonic potential. To ensure correct reorganization of the lipids and solution, the positions of all the atoms of the channel were then restrained for 2 ns. The side chains were then allowed to reorganize while the backbone was kept restrained for 8 ns. Lastly, a 100-ns unrestrained MD simulation was conducted, enabling the system to relax. The MD simulations were carried out using the GROMACS 4.6.5 program (*42*) considering a 2.0 fs time step. Bond lengths were constrained with the LINCS algorithm (*43*) and SETTLE was used for water molecules (*44*). Long-range electrostatic forces were taken into account using the particle mesh Ewald approach (*45*). A 1.2 nm cutoff was used for electrostatic and van der Waals interactions, with a switching function starting at 0.8 nm. Simulations were performed at 300 K using a Nose-Hoover thermostat (*46*). Semi-isotropic pressure control was achieved by the Parrinello-Rahman barostat (*47*). The water molecules were described using the TIP3P model (*48*). The simulation used the CHARMM22-CMAP force field with torsional cross-terms for the protein (*49*) and CHARMM36 for the phospholipids (*50*).

The simulations were performed on the BullX supercomputer Curie, TGCC, Paris, France and on superMUC, Munich, Germany with computer time awarded through the PRACE initiative.

### Multiscale modeling strategy

The activation of the voltage sensor domain in response to a depolarizing pulse is a multi-ms process resulting from a conformational transition involving several amino acids, water molecules and lipids. Developing a quantitative theory of voltage sensor activation from first principles entails the challenge of describing the time evolution of a large number of degrees of freedom over long time-scales. To tame this complexity we devised a multiscale approach, whereby enhanced sampling is achieved by biasing the equilibrium distribution of a set of auxiliary collective variables that capture the salient geometric features of the voltage sensor transition, while equilibrium properties and kinetics are investigated at a more coarse-grained level using the gating charge to characterize the free energy landscape and detect the metastable states.

### Design of Collective variables

The performance of any enhanced sampling molecular dynamics approach depends crucially on the choice of the collective variable. In the typical scenario, a biasing potential is added to the system's interactions to favor uniform sampling of the values of the collective variable. Then, if an increased value of the collective variable implies progress along the microscopic pathway, free energy barriers are easily crossed and convergence of the free energy estimation is fast. However, for many collective variables such a one-to-one correspondence between collective variable's values and intermediate states along the reaction pathway does not exist. In these cases, which we term degenerate, the degrees of freedom orthogonal to the collective variable might reach thermal equilibrium over long time-scales and the enhanced sampling approach might not be effective. One of such collective variables is the gating charge *Q*, which, despite being non-optimal for biasing purposes, is experimentally measurable and provides a transparent physical picture of the activation process. To reconcile these conflicting requirements, we have adopted a two-stage strategy whereby a free energy surface is first obtained by biasing a set of abstract CVs and is then re-computed as a function of *Q* (see Reweighting Procedure described below). This approach allowed us to achieve optimal sampling of the conformational space and, at the same time, to characterize the Gibbs free energy of experimentally relevant thermodynamic states. To enhance the sampling, we considered the two major molecular determinants of VSD activation, as determined in Delemotte et al. (*9*): (i) the unbinding [binding] of the gating charge (Ri) from [to] the initial [final] binding site and (ii) the spatial translation of the gating charges along the vector between two binding sites (Figure 2 – Figure supplement 1). Hence, progression along the activation pathway is mathematically described by expressions of the form:

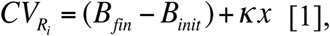

where *B*_*init*_ and *B*_*fin*_ are variables reporting the unbinding (*B*=0) or binding (*B*=1) of charge R_i_ to the initial and final binding sites, respectively; *x* is the spatial translation of the charge along the vector joining the two binding sites. *κ* makes eq. [1] dimensionally homogenous and can be used to fine-tune the relative weight of the two components. Taking advantage of the spatial proximity of several negatively charged groups of the VSD, the number of binding sites can be reduced to four: (i) phosphate groups of the top lipid layer; (ii) top protein binding site comprising E183 and E226; (iii) bottom protein binding site comprising E236 and D259; and (iv) phosphate groups of the bottom lipid bilayer (Figure 2 – Figure supplement 1). The binding collective variables *B*_*init*_ and *B*_*fin*_ depend on the coordination number *S* between the gating charges and the negative binding sites via a sigmoid function (hence the quasi-binary, on/off behavior): 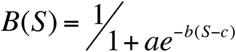, with 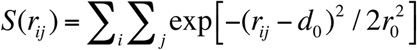, where *i* and *j* are the indices of the atoms belonging to the positively and negatively charged groups, respectively, and *r*_*ij*_ is the distance between them. The parameters *d*_*0*_, *r*_*0*_, *a, b* and *c* were chosen to fit the equilibrium distributions of bound pairs and are listed in Table 2. In this framework, each gating charge transfer can be treated separately. The transitions between the models describing the sequential states are investigated using two or three CVs. The ε*/*δ transition is described by *CV*_*R1*_ (transfer of R1 (R294) from binding sites iii to ii) and *CV*_*R3*_ (transfer of R3 (R300) from binding sites iv to iii). The δ/γ transition is described by *CV*_*R2*_ (transfer of R2 (R297) from binding sites iii to ii) and *CV*_*R4*_ (transfer of R4 (R303) from binding sites iv to iii). The γ/β transition is described by *CV*_*R1*_ (transfer of R1 (R293) from binding sites ii to i), *CV*_*R3*_ (transfer of R3 (R300) from binding sites iii to ii) and *CV*_*K5*_ (transfer of K5 (K306) from binding sites iv to iii). The β/α transition is described by *CV*_*R2*_ (transfer of R2 (R297) from binding sites ii to i), *CV*_*R4*_ (transfer of R4 (R303) from binding sites iii to ii) and *CV*_*R6*_ (transfer of R6 (R309) from binding sites iv to iii).

**Table 2.** Parameters of the switching and fitting functions used to describe the contacts between pairs of residues. The parameters for the switching function d_0_ and r_0_ were determined to fit the equilibrium distribution of distances between pairs of salt bridge partners. The fitting function is used to describe the binding in a quasi-binary fashion (a group is either bound or unbound irrespective of the number of binding partners). a, b and c are the parameters of a generalized logistic function and ensure that the distance at half-binding matches the one of the equilibrium distribution.

| | $d_0$ [nm] | $r_0$ [nm] | $a$ | $b$ | $c$ |
| --- | --- | --- | --- | --- | --- |
| R/K-$PO_4^-$ | 0.445 | 0.041 | 0.51 | 6.95 | 0.32 |
| R-E/D | 0.395 | 0.018 | 0.46 | 7.78 | 0.82 |

### Well-tempered Metadynamics

The history dependent repulsive potential added along the set of collective variables *s(x)* has the shape of a sum of Gaussians of the form:

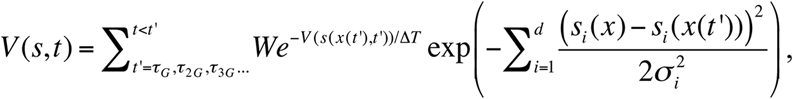

where *W* is the initial height of the Gaussian at its center, *τ* is the decrease rate of the Gaussian height along CV exploration, and *σ* is its width at half maximum height. Here, *W* is set to 0.1 kJ/mol, *ΔT* – to 30 k_B_T. *σ* is set to 0.15 for the CVs that describe the transition of a gating charge between binding sites ii and iii (CV_R1_ in ε/δ, CV_R2_ in δ/γ, CV_R3_ in γ/β and CV_R4_ in β/α) and to 0.05 for the other CVs that describe the transition of a gating charge between binding sites *Bi* and *Bii*, and *Biii* and *Biv*. Multiple walkers (42 semi-independent parallel walkers sharing the history dependent potential) metadynamics simulations were performed using MPI communication scheme implemented in the PLUMED2.1 plugin (*51*) patched to GROMACS 4.6.5 (*42*).

The use of an adaptive-bias scheme like metadynamics raises naturally a question concerning the length of the simulation: how much sampling is required to deem a free energy estimate converged? We chose to assess convergence by checking that the “instantaneous” estimates of the free energy, as obtained over the last part of the trajectories, fluctuate around a well-defined value, without any net drift (Figure 2 – Figure supplement 2). Accordingly, the errors are estimated as the standard deviation of the distribution of the estimated free energies.

### Reweighting procedure

Unlike our previous work, where we used the procedure described in Bonomi et al. (*52*), we used here a more numerically-robust procedure to calculate the time-dependent probability distribution of the conformations along the gating charge variable *Q* and thus estimate the free energy profile along this CV, as described in Tiwary et al. (*53*). The weighted histogram along *Q* is built using 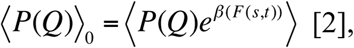 where *F(s,t)* is the time- and position-dependent estimate of the free energy:

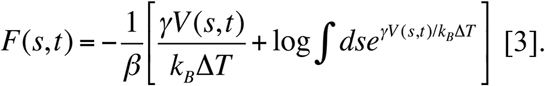

Here, *V(s,t)* is the time- and position-dependent bias, and *ΔT* is the decrease rate of the Gaussian height of the well-tempered metadynamics run.

Despite the fact that this reweighting procedure is mathematically rigorous, it nevertheless relies on the assumption of instantaneous equilibrium (*53*). To estimate the accuracy of the reweighting procedure and evaluate the portion of the trajectory to be discarded, we have compared the free energy estimated along the original CVs using the direct estimator [3] and passing through the reweighting step [2] for each of the transition, using the entire dataset, the last half or the last quarter of the trajectories (Figure 2 – Figure supplement 4-7). For the two 2D-transitions, we have considered only the last half of the 10 μs simulation as on this subset of configuration different estimators for the free energy provide consistent results (with discrepancies of the order of 0.5 kcal/mol, see Figure 2 – Figure supplement 6-7). For the two three-dimensional transitions, on the other hand, the volume of the configurational space to sample in order to achieve convergence is significantly larger; therefore we considered the entire dataset of configurations. The estimated error is then ~ 2 kcal/mol (Figure 2 – Figure supplement 4-5).

### Gating charge expression

We used the trajectories sampled along the metadynamics runs to compute the potential of mean force (PMF) as a function of the gating charge *Q*. *Q* can be linked to the microscopic state of the channel through: 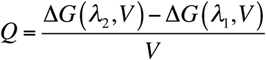, where *V* is the TM potential. For each channel conformation (*λ*, being a set of atomic coordinates), Δ*G*(*λ*,*V*) is the reversible work component due to the applied voltage *V*. It relates the conformation of the channel *λ* to 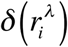, the so-called “electrical” distance (*54*): 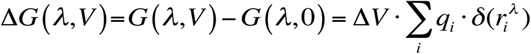, where *q*_*i*_ is the *i*^*th*^ - protein charge and 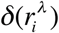 is the “electrical distance”, given by: 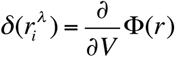. This quantity accounts for the degree of coupling between the local electrostatic potential at the position of charge *q*_*i*_, Φ(*r*), and the TM applied potential, *V*. In practice, 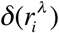 is evaluated for each protein configuration *λ* by carrying out two independent simulations of the system under two different TM potentials *V* (*55*). For each *V*, the local-electrostatic potential Φ(***r***) is then calculated as an average over n=100 configurations sampled along 2 ns of simulation. For a given conformation *λ*, the electrical distances are estimated as *δ* ≡ [*φ*^λ^ (**r**,Δ*V*_2_) *– φ* ^*λ*^(**r**,Δ*V*_1_)]/(Δ*V*_2_ – Δ*V*_1_). Here, based on our previous study (*9*), we used the facts that (i) *Q* can be associated to each configuration by summing the contributions from all the charged residues located in the narrowest region of the VSD, i.e. where the electrostatic potential changes abruptly, and (ii) 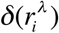 can be very well approximated by the best fit of the measured electrical distance in the different states by the generalized logistic function 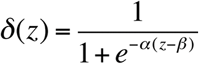 depending only on the z position of the charged residues of the VSD, with *α* = 1.80654 and *β* = 0.0949.

### Gating current calculations

Simulating gating charge kinetics based on the shape of the PMF required an assumption of governing dynamics. The large friction approximation yields the Smoluchowski equation, which was solved using a variety of methods as summarized in the text. The methods have been previously described (*1,36*).

The prefactor *D* was initially estimated as the diffusion coefficient of the gating charge collective variable *Q*. For the four transitions, the bin size considered was 0.2 e. The diffusion coefficient of the gating charge was estimated on a 40 ns MD simulation performed on a flattened free energy landscape via its mean square displacement (MSD): 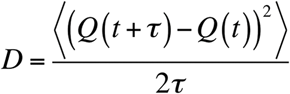, where *τ* is the time lag. In particular, the slope of the MSD along *τ* was calculated during the interval between 400 and 4000 ps.

### Diffusion maps

Since in the multiple-walker scheme the exploration is initiated from several distinct structures in the ε, δ, γ, β and α states, a crucial aspect to consider is whether or not the set of configurations sampled along the trajectory span a connected domain of the configurational space and thus if relative weights can be always correctly assigned. To test this hypothesis, we used a simple criterion from spectral graph theory, namely the absence of gaps in the spectrum of an appropriately defined discrete Laplace operator (*56*). The latter is defined in terms of an adjacency matrix built from pairwise root mean square values after a Gaussian kernel of bandwidth *σ* has been applied (Figure 2 – Figure supplement 7).

We estimated *ε* as 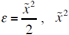 being the median of the rmsd between the closest 0.5 % configurations. It amounts to 4.2 Å for the configurations taken from the first tenth of the ensemble of the well-tempered metadynamics trajectories and to 9.0 Å for the ensemble made of configuration from the entire metadynamics trajectories. When computed over the first tenth of the ensemble of the well-tempered metadynamics trajectories, the eigenvalue spectrum is features a gap after the third eigenvalue. Over the ensemble of entire trajectories, on the other hand, the eigenvalue spectrum decreases smoothly and regularly, indicating that the configurational ensemble has been populated beyond the starting configurations ε, δ, γ, β and α.

## Acknowledgements

We acknowledge PRACE for awarding us access to resource CURIE based in France at the TGCC and SUPERMUC based in Germany at the LRZ. Preliminary results were obtained thanks to generous allocation of computer time from GENCI France (ID-panel-year). This work was supported in part by National Institutes of Health Grant National Institute of General Medical Sciences (NIGMS) P01 55876 (to M.L.K) and the Commonwealth of Pennsylvania. LD receives funding from the European Union Framework Program (PIOF-GA-2012-329534) “Voltsens.” We are grateful to Fabrizio Marrinelli for fruitful discussions.

## Statement

We have no conflicts of interest to declare.

---

**Figure 1 – Figure Supplement 1.**
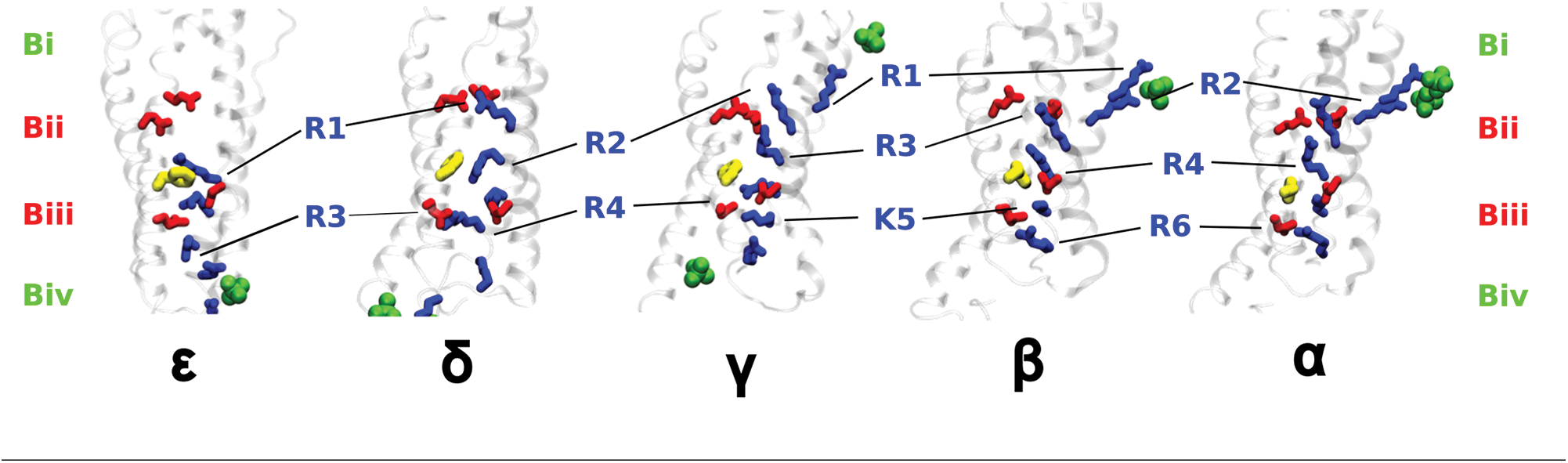
Structures of the five initial states, ranging from the most resting (ε) to the most activated (α). State-dependent salt bridge pairs between positively charged residues of S4 (blue) and negative charges from the protein (red) or the lipids (green) are shown. The negative charges are grouped into four binding sites (Bi to Biv). F233 is represented in yellow. The two or three positively charged residues, which are subject to the bias potential, are labeled in blue.

**Figure 2 – Figure Supplement 1.**
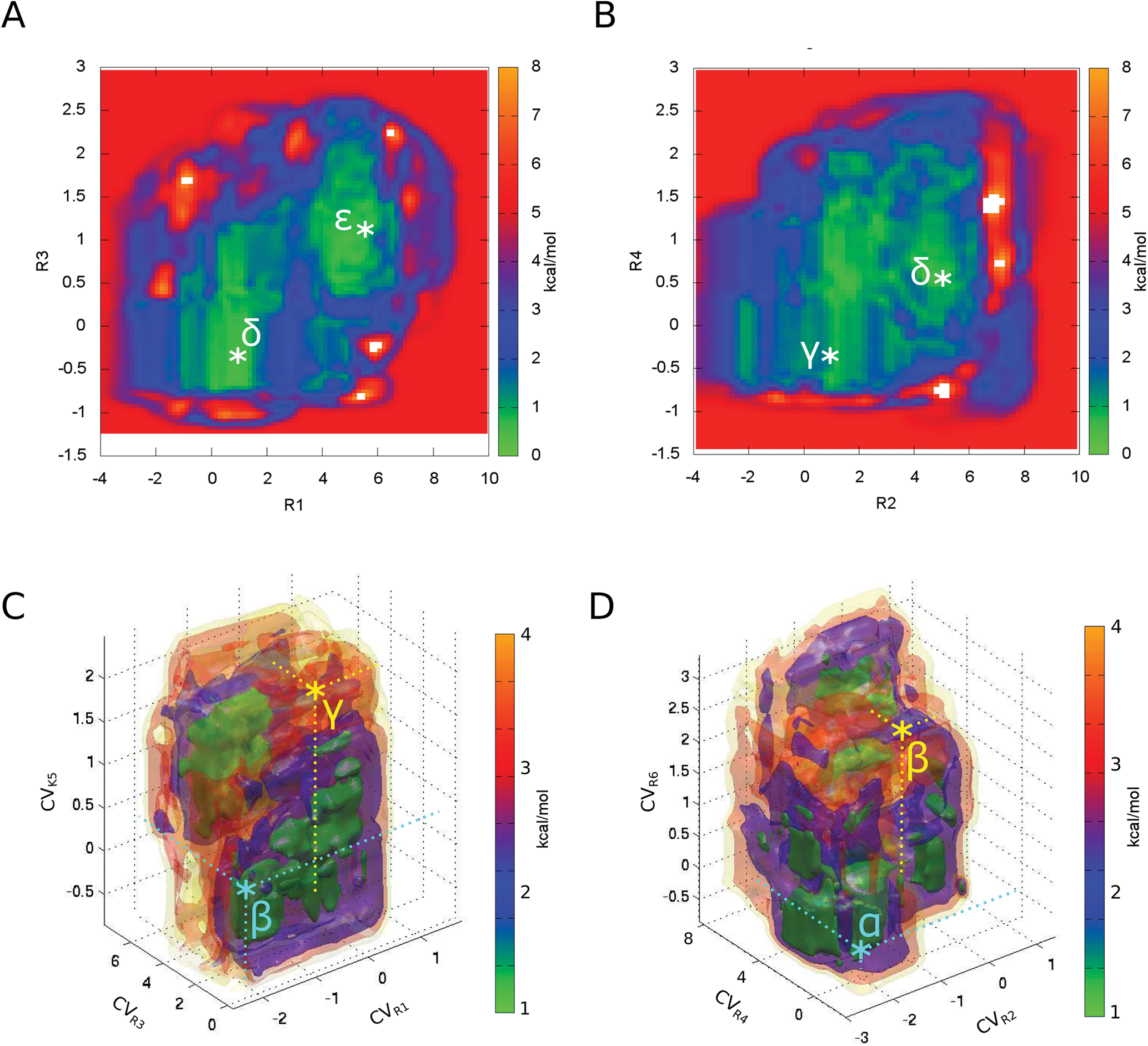
Standard deviation from the average free energy in the original extended geometrical collective variable space (as shown in Figure 2B of the Main Text), as computed over the last half of the trajectories. A, B, C and D correspond respectively to the ε/δ, δ/γ, γ/β and β/a transitions. The initial configurations are highlighted with the * symbol.

**Figure 2 – Figure Supplement 2.**
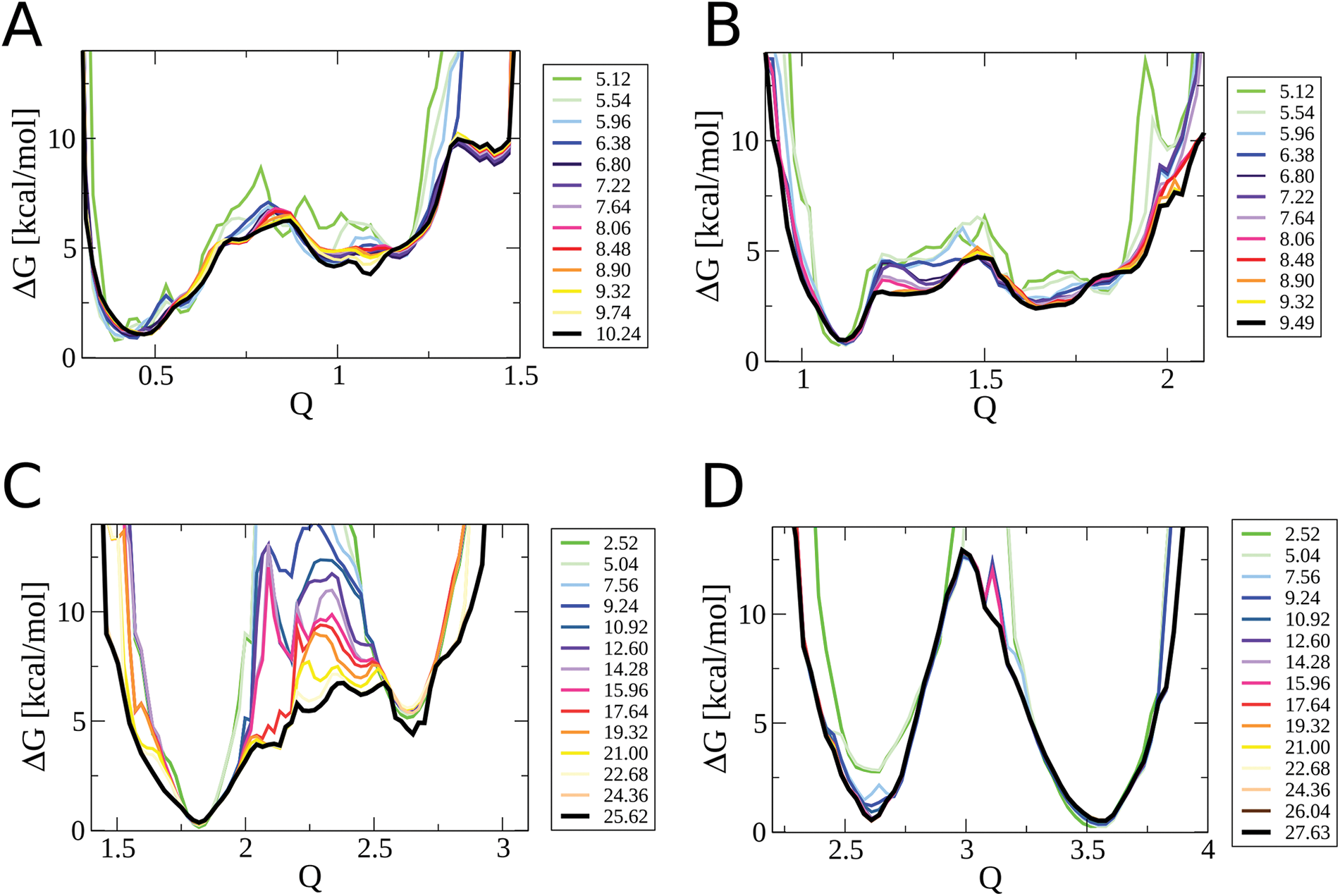
Time (in µs) evolution of the PMF along the gating charge (Q in units of eu) collective variable for the four separate transitions. A, B, C and D correspond to the E/Δ, Δ/Γ, Γ/B and B/A transitions respectively.

**Figure 2 – Figure Supplement 3.**
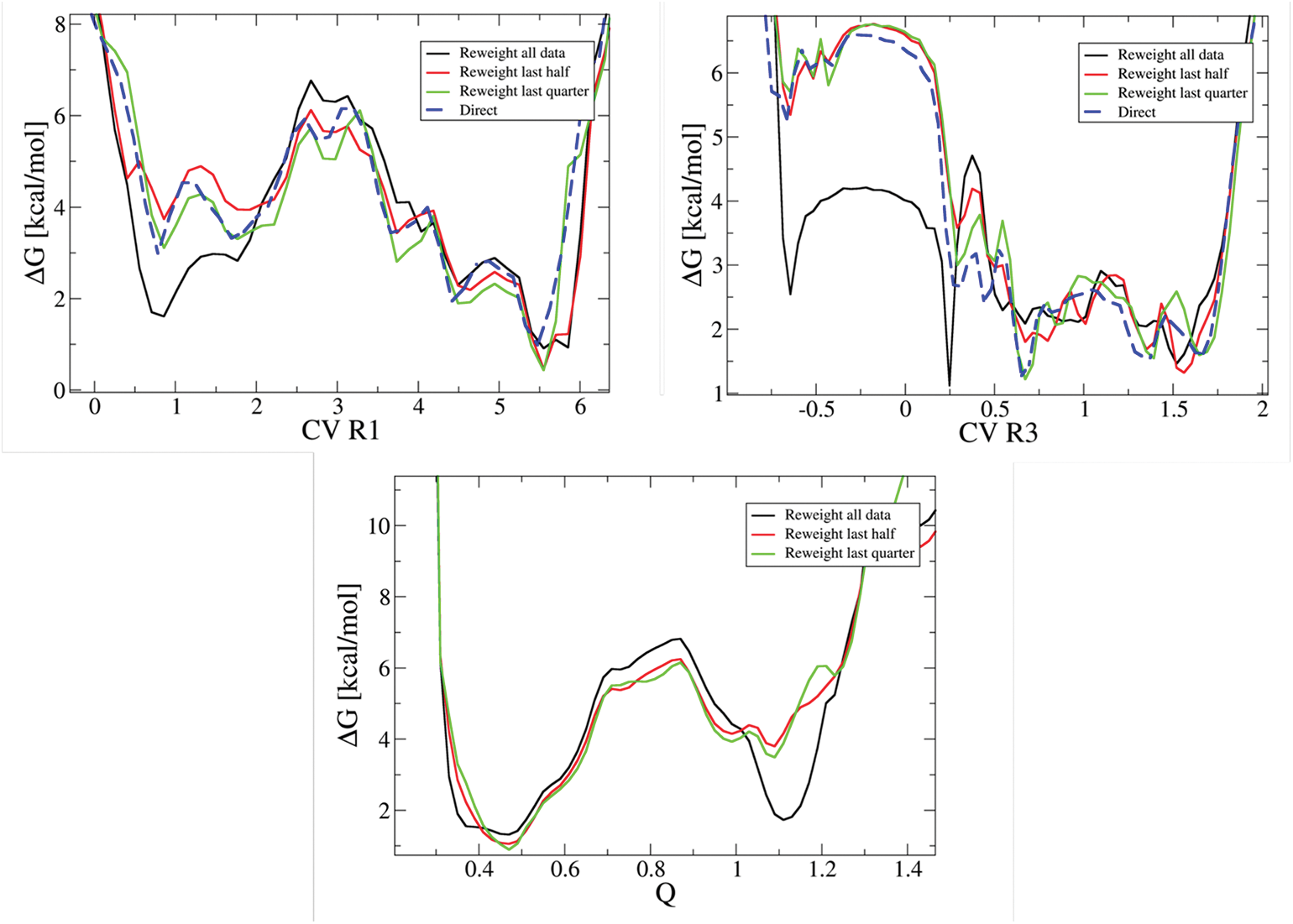
Comparison between the Ε/Δ PMF in the original extended geometrical collective variable space (CV_R1_ and CV_R3_) and in the reweighted collective variable space (Q in units of eu), when measured directly from the metadynamics repulsive potential (blue) or after reweighting the entire data set (black), the last half (red) or the last quarter (green).

**Figure 2 – Figure Supplement 4.**
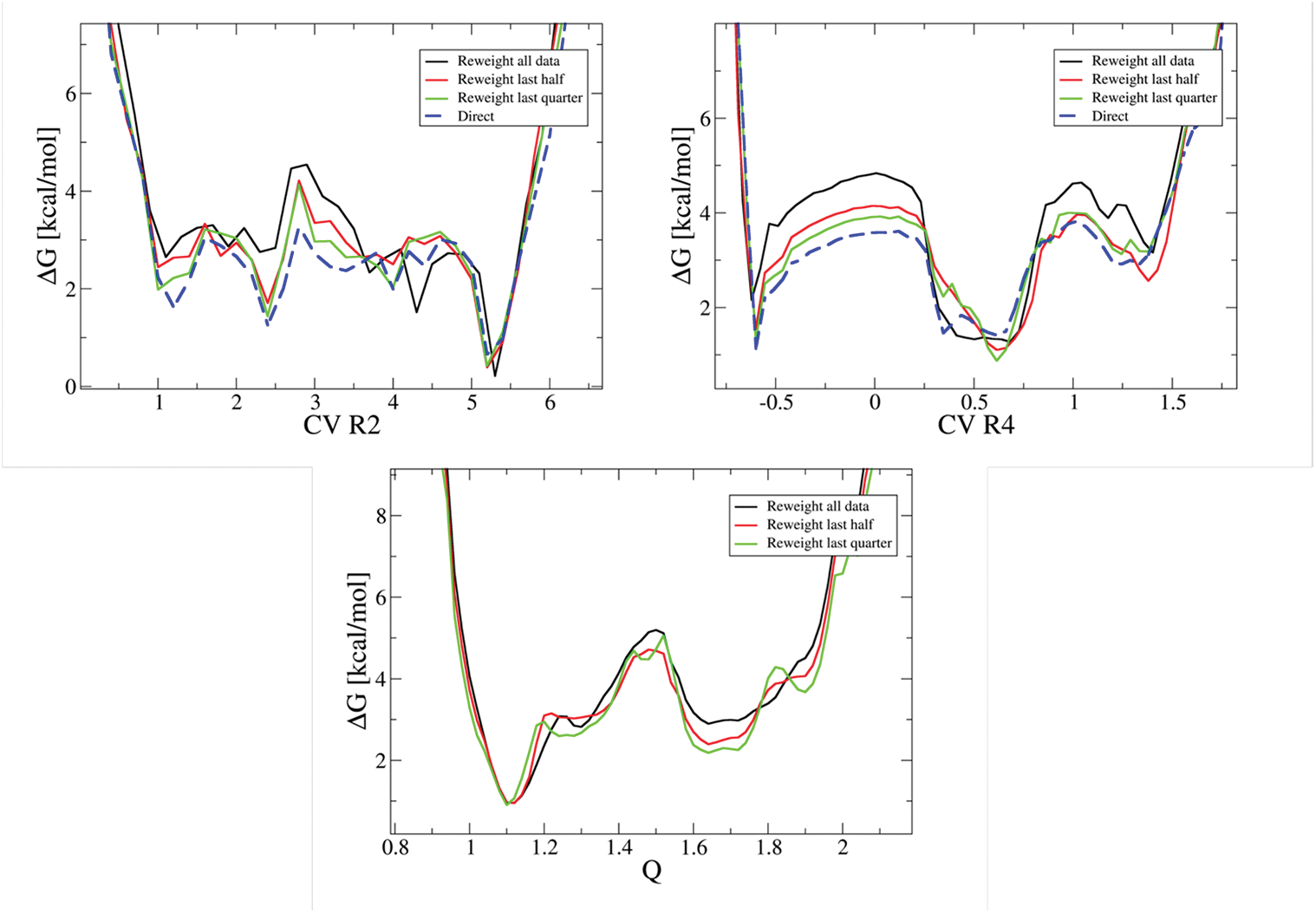
Comparison between the Δ/Γ PMF in the original extended geometrical collective variable space (CV_R2_ and CV_R4_) and in the reweighted collective variable space (Q in units of eu), when measured directly from the metadynamics repulsive potential (blue) or after reweighting the entire data set (black), the last half (red) or the last quarter (green).

**Figure 2 – Figure Supplement 5.**
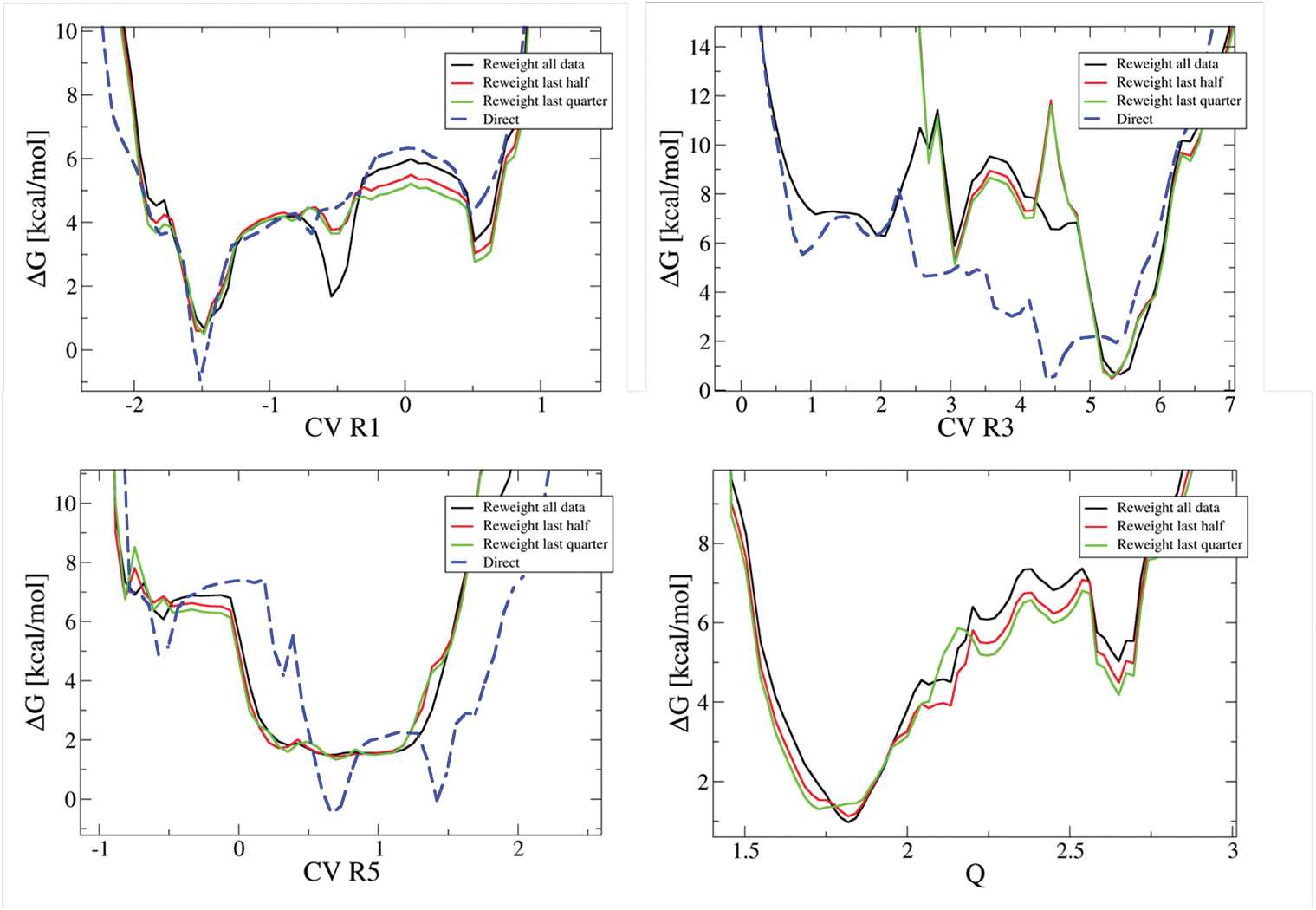
Comparison between the Γ/B PMF in the original extended geometrical collective variable space (CV_R1_, CV_R3_ and CV_K5_) and in the reweighted collective variable space (Q in units of eu), when measured directly from the metadynamics repulsive potential (blue) or after reweighting the entire data set (black), the last half (red) or the last quarter (green).

**Figure 2 – Figure Supplement 6.**
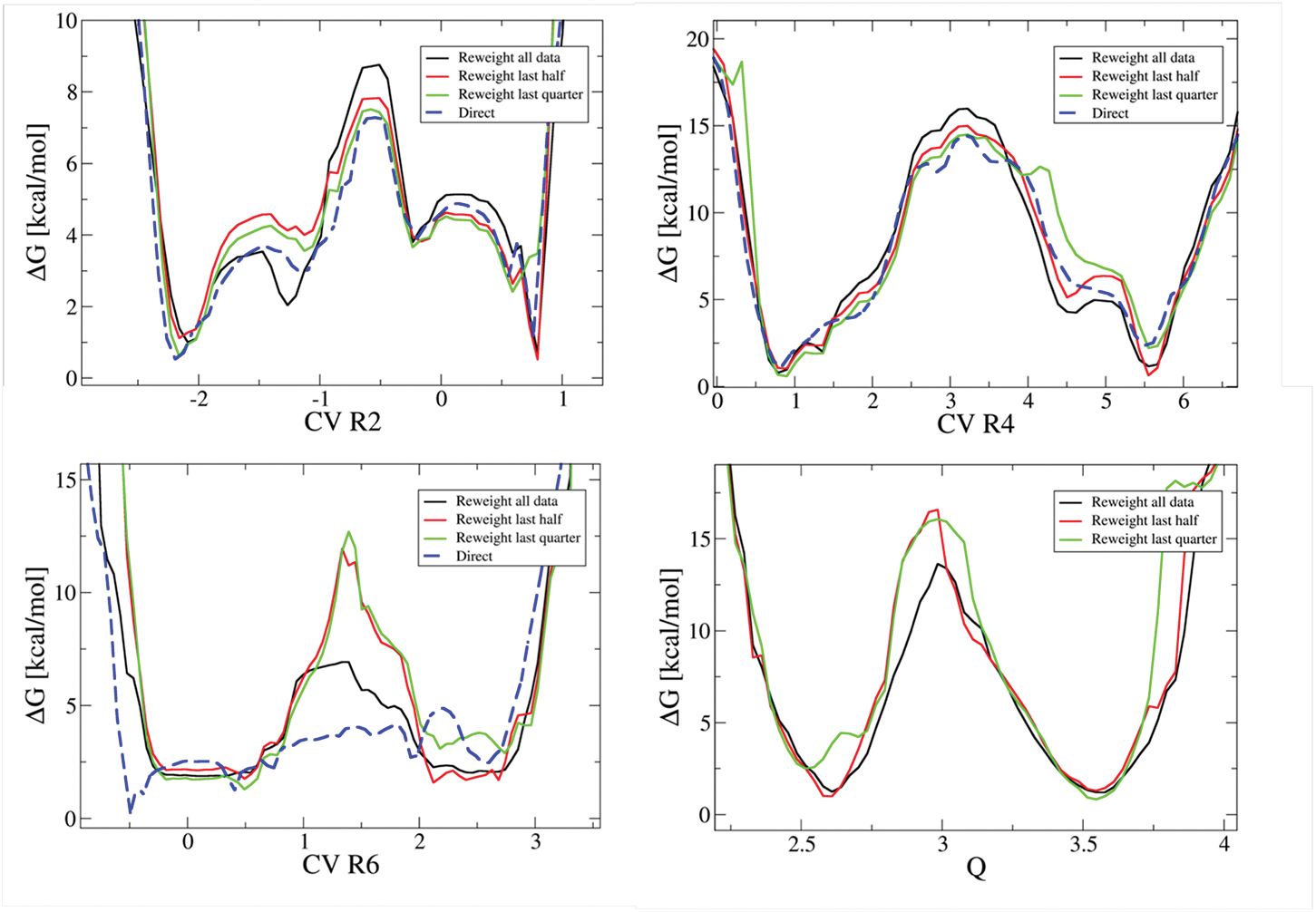
Comparison between the B/A PMF in the original extended geometrical collective variable space (CV_R2_, CV_R4_ and CV_R6_) and in the reweighted collective variable space (Q in units of eu), when measured directly from the metadynamics repulsive potential (blue) or after reweighting the entire data set (black), the last half (red) or the last quarter (green).

**Figure 2 – Figure supplement 7.**
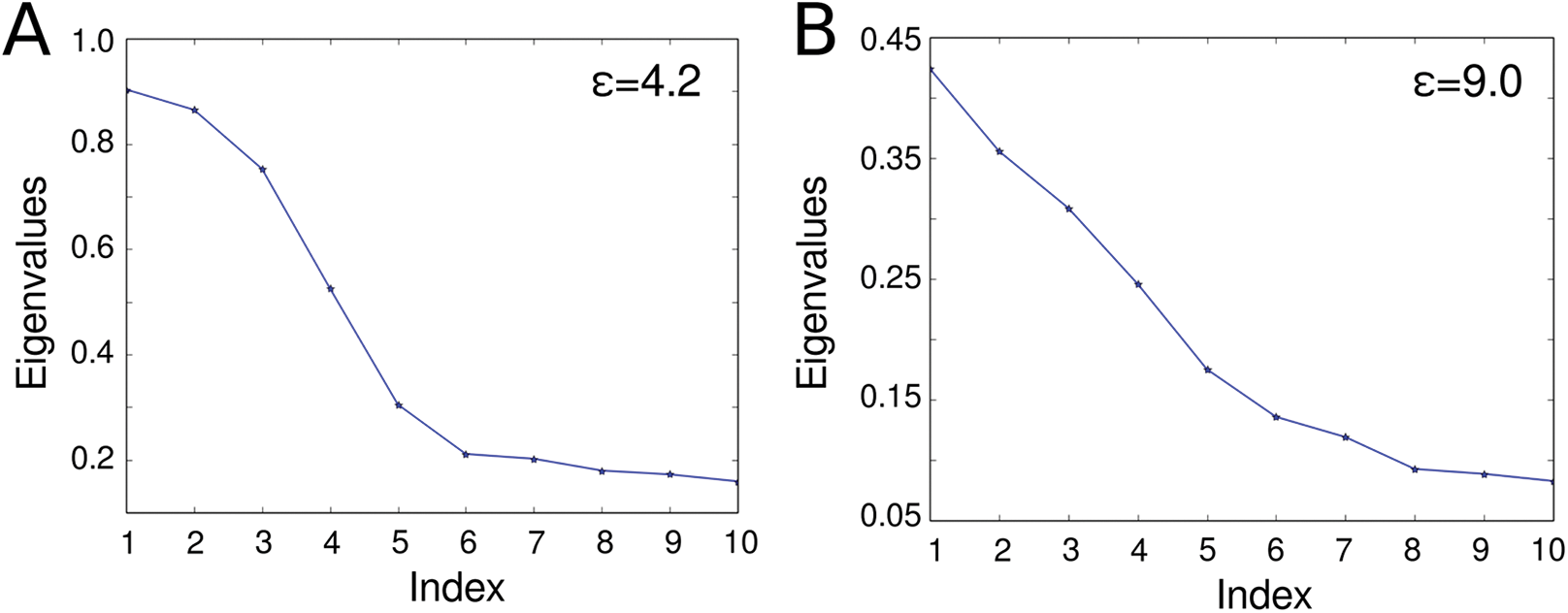
Connectivity of the five states. Eigenvalues spectrum of a discrete Laplace operator, defined in terms of an adjacency matrix built from pairwise root mean square values after a Gaussian kernel of bandwidth ε has been applied, associated with the first 10% of the concatenated trajectory (A) and the entire concatenated trajectory (B).

## Supplement Movie 1

Reconstructed deactivation trajectory, highlighting the rearrangement of the voltage sensor structural elements. The ~ 1500 snapshots extracted from the ensemble of configurations along the metadynamics runs were ordered decreasing values of the gating charge Q. The movie was generated using VMD, considering a trajectory smoothing window size of 1. The colors of the different structural elements of the voltage sensor are described in (Figure 3 – Figure Supplement 2)

**Figure 3 – Figure Supplement 1.**
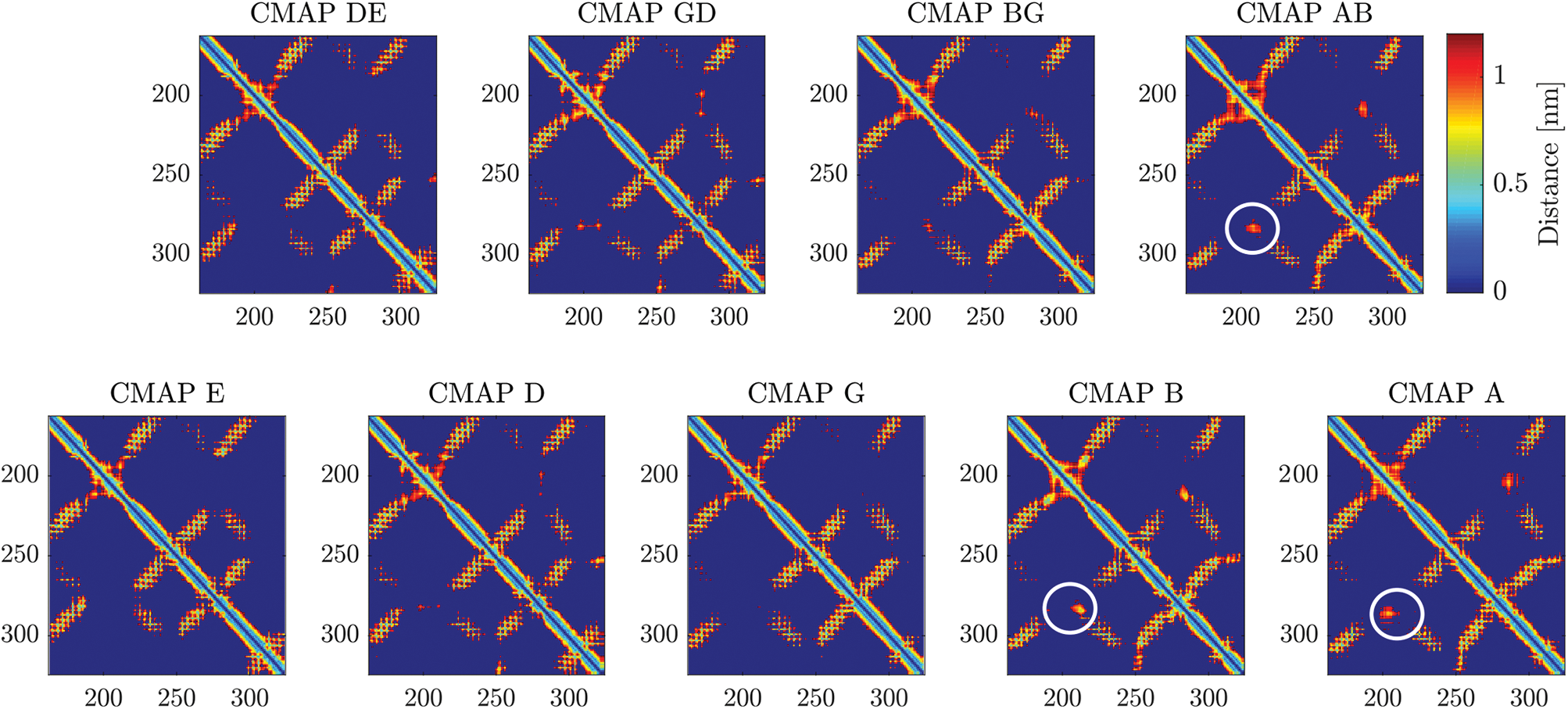
Contact maps between pairs of residues in the different activation and transition states. The minimum distance between the two residues is reported (in nm). Distances beyond 1.2 nm are shown in dark blue. The interaction between the S1-S2 and the S3-S4 linkers, highlighted with white circles, is significantly reshaped during the B/A transition.

**Figure 3 – Figure Supplement 2.**
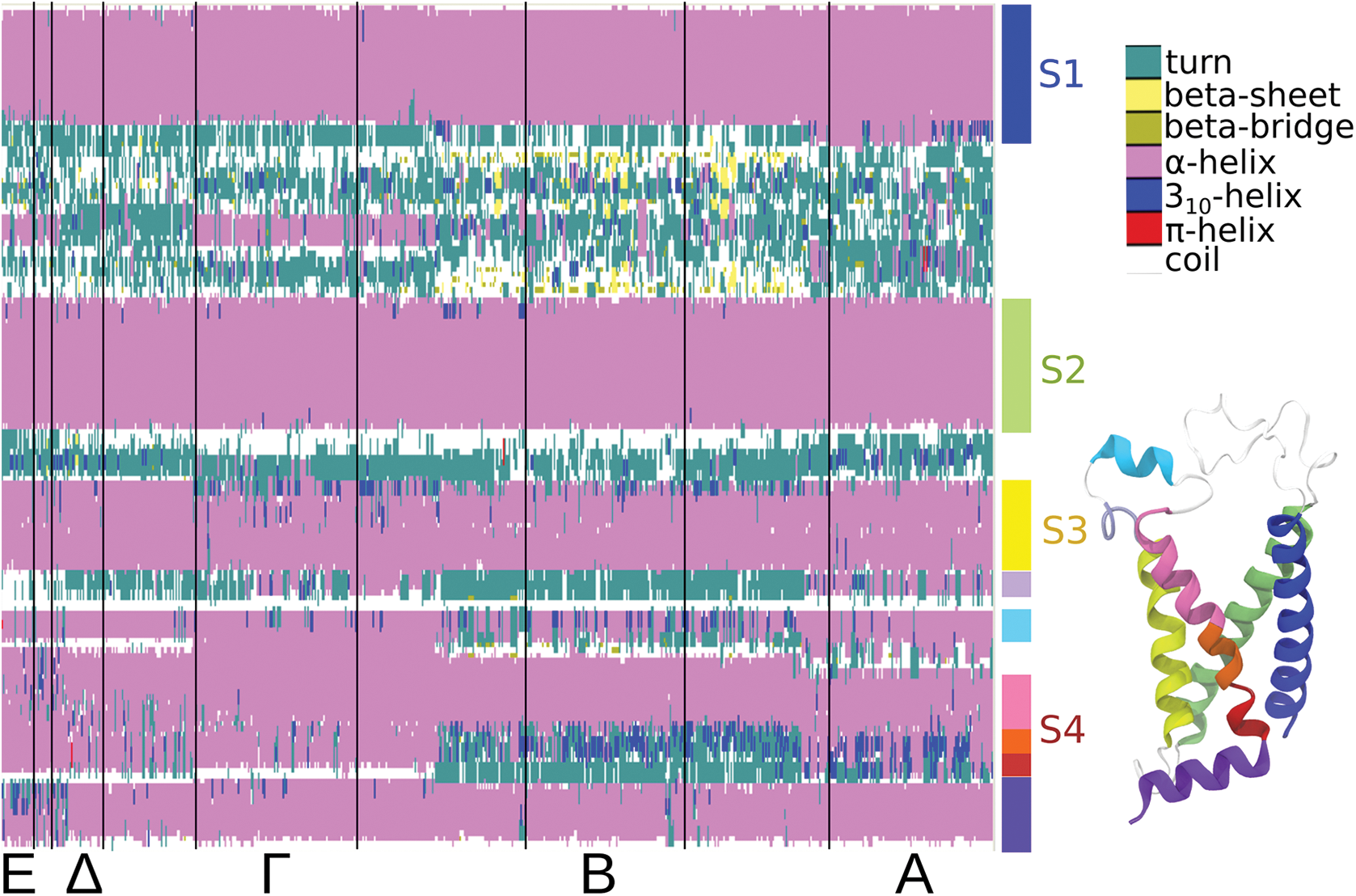
Per-residue secondary structure map estimated along the activation pathway (horizontal axis) obtained using the Timeline plugin for VMD (*32*). The vertical axis corresponds to the sequence of the voltage sensor domain. Along the horizontal axis, the configurations are ordered according to increasing values of Q; the vertical lines separate the configurations belonging to the E, Δ, Γ, B and A ensembles. Different structural elements of the voltage sensor are shown as colored boxes to the left of the secondary structure map and also represented as ribbons of the same color on the molecular structure in the right bottom corner.

**Figure 7 – Figure supplement 1.**
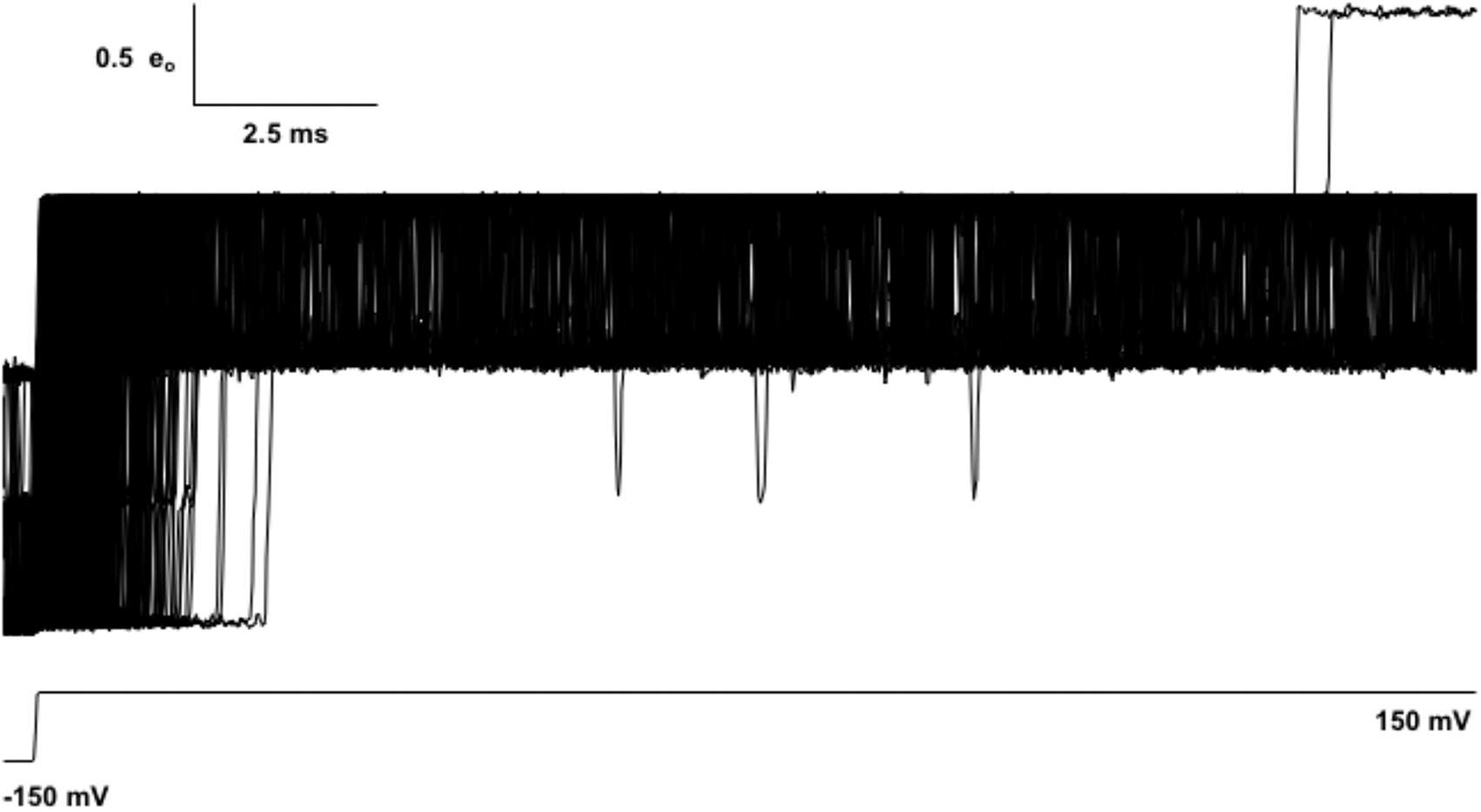
1000 Brownian motion trajectories of gating charge Q in response to a test pulse to +150 mV from a holding potential V_h_ = −150 mV. The A state is reached only twice despite the large forward bias in voltage.

## References and Notes

1. D. Sigg, Modeling ion channels: Past, present, and future. J. Gen. Physiol. 144, 7–26 (2014).

2. T. Andersson, Exploring voltage-dependent ion channels in silico by hysteretic conductance. Math. Biosci. 226, 16–27 (2010).

3. X. Tao, A. Lee, W. Limapichat, D. A. Dougherty, R. MacKinnon, A gating charge transfer center in voltage sensors. Science. 328, 67–73 (2010).

4. U. Henrion, J. Renhorn, S. I. Börjesson, E. M. Nelson, C. S. Schwaiger, P. Bjelkmar, B. Wallner, E. Lindahl, F. Elinder From the Cover: Tracking a complete voltage-sensor cycle with metal-ion bridges. Proc. Natl. Acad. Sci. 109, 8552–8557 (2012).

5. D. Wu, K. Delaloye, M. A. Zaydman, A. Nekouzadeh, Y. Rudy, J. Cui, State-dependent electrostatic interactions of S4 arginines with E1 in S2 during Kv7.1 activation. J Gen Physiol. 135, 595–606 (2010).

6. P. G. DeCaen, V. Yarov-Yarovoy, E. M. Sharp, T. Scheuer, W. A. Catterall, Sequential formation of ion pairs during activation of a sodium channel voltage sensor. Proc Acad Natl Sci USA. 106, 22498 (2009).

7. B. S. Long, E. B. Campbell, R. Mackinnon, Voltage sensor of Kv1.2: Structural basis of electromechanical coupling. Science. 309, 903–908 (2005).

8. J. Payandeh, T. Scheuer, N. Zheng, W. A. Catterall, The crystal structure of a voltage-gated sodium channel. Nature. 475, 353–358 (2011).

9. L. Delemotte, M. Tarek, M. L. Klein, C. Amaral, W. Treptow, Intermediate states of the Kv1.2 voltage sensor from atomistic molecular dynamics simulations. Proc Acad Natl Sci USA. 108, 6109–6114 (2011).

10. E. Vargas, V. Yarov-Yarovoy, F. Khalili-Araghi, W. A. Catterall, M. L. Klein, M. Tarek, E. Lindahl, K. Schulten, E. Perozo, F. Bezanilla, B. Roux An emerging consensus on voltage-dependent gating from computational modeling and molecular dynamics simulations. J. Gen. Physiol. 140, 587–594 (2012).

11. J. A. Freites, E. V. Schow, S. H. White, D. J. Tobias, Microscopic origin of gating current fluctuations in a potassium channel voltage-sensor. Biophys J. 102, A44–A46 (2012).

12. M. Ø. Jensen, V. Jogini, D. avi. W. Borhani, A. E. Leffler, R. O. Dror, D. E. Shaw, Mechanism of Voltage Gating in Potassium Channels. Science. 336, 229–233 (2012).

13. F. Khalili-Araghi, V. Jogini, V. Yarov-Yarovoy, E. Tajkhorshid, B. Roux, K. Schulten, Calculation of the gating charge for the Kv1.2 voltage activated potassium channel. Biophys J. 98, 1–10 (2010).

14. J. R. Silva, H. Pan, D. Wu, A. Nekouzadeh, K. F. Decker, J. Cui, N. a. Baker, D. Sept, Y. Rudy, A multiscale model linking ion-channel molecular dynamics and electrostatics to the cardiac action potential. Proc. Natl. Acad. Sci. 106, 11102–11106 (2009)

15. C. S. Souza, C. Amaral, W. Treptow, Electric fingerprint of voltage sensor domains. Proc. Natl. Acad. Sci. 111, 17510–17515 (2014).

16. E. Palovcak, L. Delemotte, M. L. Klein, V. Carnevale, Evolutionary imprint of activation: The design principles of VSDs. J. Gen. Physiol. 143, 145–156 (2014).

17. I. S. Ramsey, M. M. Moran, J. A. Chong, D. E. Clapham, A voltage-gated proton-selective channel lacking the pore domain. Nature. 440, 1213–1216 (2006).

18. Y. Murata, H. Iwasaki, M. Sasaki, K. Inaba, Y. Okamura, Phosphoinositide phosphatase activity coupled to an intrinsic voltage sensor. Nature. 435, 1239–1243 (2005).

19. J. Zhao, R. Blunck, The isolated voltage sensing domain of the Shaker potassium channel forms a voltage-gated cation channel. eLife. 5, e18130 (2016).

20. G. A. Haddad, R. Blunck, Mode shift of the voltage sensors in Shaker K+ channels is caused by energetic coupling to the pore domain. J. Gen. Physiol. 137, 455–472 (2011).

21. P. S. Tan, M. D. Perry, C. A. Ng, J. I. Vandenberg, A. P. Hill, Voltage-sensing domain mode shift is coupled to the activation gate by the N-terminal tail of hERG channels. J. Gen. Physiol. 140, 293–306 (2012).

22. A. Bruening-Wright, H. P. Larsson, Slow Conformational Changes of the Voltage Sensor during the Mode Shift in Hyperpolarization-Activated Cyclic-Nucleotide-Gated Channels. J. Neurosci. 27, 270–278 (2007).

23. C. A. Villalba-Galea, W. Sandtner, D. M. Starace, F. Bezanilla, S4-based voltage sensors have three major conformations. Proc. Natl. Acad. Sci. 105, 17600 (2008).

24. R. Olcese, R. Latorre, L. Toro, F. Bezanilla, E. Stefani, Correlation between Charge Movement and Ionic Current during Slow Inactivation in Shaker K+ Channels. J. Gen. Physiol. 110, 579–589 (1997).

25. R. Männikkö, S. Pandey, H. P. Larsson, F. Elinder, Hysteresis in the Voltage Dependence of HCN Channels. J. Gen. Physiol. 125, 305–326 (2005).

26. J. Guo, W. Zeng, Q. Chen, C. Lee, L. Chen, Y. Yang, C. Cang, D. Ren, Y. Jiang, Structure of the voltage-gated two-pore channel TPC1 from Arabidopsis thaliana. Nature. 531, 196–201 (2016).

27. Q. Li, S. Wanderling, M. Paduch, D. Medovoy, A. Singharoy, R. McGreevy, C. A. Villalba-Galea, R. E. Hulse, B. Roux, K. Schulten, A. Kossiakoff, E. Perozo, Structural mechanism of voltage-dependent gating in an isolated voltage-sensing domain. Nat. Struct. Mol. Biol. 21, 244–252 (2014).

28. B. S. Long, E. B. Campbell, R. MacKinnon, Crystal structure of a mammalian voltage-dependent Shaker family K+ channel. Science. 309, 897–903 (2005).

29. L. Delemotte, M. A. Kasimova, M. L. Klein, M. Tarek, V. Carnevale, Free-energy landscape of ion-channel voltage-sensor–domain activation. Proc. Natl. Acad. Sci. 112, 124–129 (2015).

30. A. Laio, F. L. Gervasio, Metadynamics: a method to simulate rare events and reconstruct the free energy in biophysics, chemistry and material science. Rep Prog Phys. 71, 126601 (2008).

31. C. S. Schwaiger, P. Bjelkmar, B. Hess, E. Lindahl, 310-helix conformation facilitates the transition of a voltage sensor S4 segment toward the down state. Biophys J. 100, 1446–1454 (2011).

32. W. Humphrey, A. Dalke, K. Schulten, VMD - Visual Molecular Dynamics. J Molec Graph. 14, 33–38 (1996).

33. I. G. Ishida, G. E. Rangel-Yescas, J. Carrasco-Zanini, L. D. Islas, Voltage-dependent gating and gating charge measurements in the Kv1.2 potassium channel. J. Gen. Physiol. 145, 345–358 (2015).

34. D. Sigg, F. Bezanilla, E. Stefani, Fast gating in the Shaker K+ channel and the energy landscape of activation. Proc Nat Acad Sci USA. 100, 7611 (2003).

35. D. Sigg, H. Qian, F. Bezanilla, Kramers’ diffusion theory applied to gating kinetics of voltage-dependent ion channels. Biophys. J. 76, 782–803 (1999).

36. D. Sigg, Gating current noise produced by elementary transitions in Shaker potassium channels. Science. 264, 578 (1994).

37. W. N. Zagotta, T. Hoshi, R. W. Aldrich, Shaker potassium channel gating. III: Evaluation of kinetic models for activation. J Gen Physiol. 103, 321 (1994).

38. B. M. Rodríguez, D. Sigg, F. Bezanilla, Voltage Gating of Shaker K+ Channels The Effect of Temperature on Ionic and Gating Currents. J. Gen. Physiol. 112, 223–242 (1998).

39. M. F. Priest, J. J. Lacroix, C. A. Villalba-Galea, F. Bezanilla, S3-S4 Linker Length Modulates the Relaxed State of a Voltage-Gated Potassium Channel. Biophys. J. 105, 2312–2322 (2013).

40. X. Chen, Q. Wang, F. Ni, J. Ma, Structure of the full-length Shaker potassium channel Kv1. 2 by normal-mode-based X-ray crystallographic refinement. Proc Acad Natl Sci USA. 107, 11352 (2010).

41. A. Sali and T. L. Blundell. Comparative protein modelling by satisfaction of spatial restraints. J. Mol. Biol. 234, 779–815, 1993.

42. B. Hess, C. Kutzner, D. van der Spoel, E. Lindahl, GROMACS 4: algorithms for highly efficient, load-balanced, and scalable molecular simulation. J Comp Theor Chem. 4, 435–447 (2008).

43. B. Hess, H. Bekker, H. J. C. Berendsen, J. G. E. M. Fraaije, LINCS: A linear constraint solver for molecular simulations. J. Comput. Chem. 18, 1463–1472 (1997).

44. S. Miyamoto, P. A. Kollman, Settle: An analytical version of the SHAKE and RATTLE algorithm for rigid water models. J. Comput. Chem. 13, 952–962 (1992).

45. T. Darden, D. York, L. Pedersen, Particle Mesh Ewald - an Nlog(N) method for ewald sums in large systems. J Chem Phys. 98, 10089–10092 (1993).

46. S. Nosé, A unified formulation of the constant temperature molecular dynamics methods. J Chem Phys. 81, 511–519 (1984).

47. M. Parrinello, A. Rahman, Polymorphic transitions in single crystals: A new molecular dynamics method. J. Appl. Phys. 52, 7182–7190 (1981).

48. W. L. Jorgensen, J. Chandrasekhar, J. D. Madura, R. W. Impey, M. L. Klein, Comparison of simple potential functions for simulating liquid water. J Chem Phys. 79, 926–935 (1983).

49. MacKerell, M. Feig, C. L. Brooks, Improved Treatment of the Protein Backbone in Empirical Force Fields. J. Am. Chem. Soc. 126, 698–699 (2004).

50. J. B. Klauda, R. M. Venable, J. A. Freites, J. W. O’Connor, D. J. Tobias, C. Mondragon-Ramirez, I. Vorobyov, A. D. MacKerell, R. W. Pastor, Update of the CHARMM All-Atom Additive Force Field for Lipids: Validation on Six Lipid Types. J Phys Chem B. 114, 7830–7843 (2010).

51. G. A. Tribello, M. Bonomi, D. Branduardi, C. Camilloni, G. Bussi, PLUMED 2: New feathers for an old bird. Comput. Phys. Commun. 185, 604–613 (2014).

52. M. Bonomi, A. Barducci, M. Parrinello, Reconstructing the equilibrium Boltzmann distribution from well-tempered metadynamics. J. Comput. Chem. 30, 1615–1621 (2009).

53. P. Tiwary, M. Parrinello, A Time-Independent Free Energy Estimator for Metadynamics. J. Phys. Chem. B. 119, 736–742 (2015).

54. F. J. Sigworth, Voltage gating of ion channels. Q Rev Biophys. 27, 1–40 (1994).

55. W. Treptow, M. Tarek, M. L. Klein, Initial Response of the Potassium Channel Voltage Sensor to a Transmembrane Potential. J. Am. Chem. Soc. 131, 2107–2109 (2009).

56. R. R. Coifman, S. Lafon, a B. Lee, M. Maggioni, B. Nadler, F. Warner, S. W. Zucker, Geometric diffusions as a tool for harmonic analysis and structure definition of data: Diffusion maps. Proc. Natl. Acad. Sci. U. S. A. 102, 7426–7431 (2005).

